# A Systematic Interrogation of MHC Class I Peptide Presentation Identifies Constitutive and Compensatory Protein Degradation Pathways

**DOI:** 10.1101/2021.10.07.463289

**Authors:** Jennifer L. Mamrosh, David J. Sherman, Joseph R. Cohen, James A. Johnston, Marisa K. Joubert, Jing Li, J. Russell Lipford, Brett Lomenick, Annie Moradian, Siddharth Prabhu, Michael J. Sweredoski, Bryan Vander Lugt, Rati Verma, Raymond J. Deshaies

## Abstract

The adaptive immune system distinguishes self from non-self by surveying peptides generated from degradation of intracellular proteins that are loaded onto MHC Class I molecules for display on the cell surface. While early studies reported that the bulk of cell surface MHC Class I complexes require the ubiquitin-proteasome system (UPS) for their generation, this conclusion has been challenged. To better understand MHC Class I peptide origins, we sought to carry out a comprehensive, quantitative census of the MHC Class I peptide repertoire in the presence and absence of UPS activity. We introduce optimized methodology to enrich for authentic Class I-bound peptides in silico and then quantify by mass spectrometry their relative amounts upon perturbation of the ubiquitin-proteasome system. Whereas most peptides are dependent on the proteasome and ubiquitination for their generation, a surprising 30% of the MHC Class I repertoire, enriched in peptides of mitochondrial origin, appears independent of these pathways. A further ∼10% of Class I-bound peptides were found to be dependent on the proteasome but independent of ubiquitination for their generation. Notably, clinically achievable partial inhibition of the proteasome resulted in display of novel peptides antigens, at least one of which promotes immune system activation. Our results suggest that generation of MHC Class I•peptide complexes is more complex than previously recognized and also provide evidence for compensatory peptide-generating pathways when canonical pathways are impaired.

## INTRODUCTION

To survive, organisms must be able to detect threats in their environment. While all multicellular organisms have an innate immune system broadly surveying for common pathogens, evolution of an adaptive immune system allowed for specific and lasting responses to wider threats. Central to adaptive immunity is the display of antigens. All nucleated cells in jawed vertebrates display on their cell surface peptide antigens derived from intracellular proteins. This represents the current state of cells to the immune system and allows for the detection of intracellular infection by bacteria and viruses. Peptides typically nine amino acids in length, generated from intracellular proteins, are presented in a non-covalent complex with the plasma membrane protein MHC Class I. These peptides, largely originating as longer peptides generated by proteasomal degradation, often are subject to additional trimming by proteases in the cytoplasm and endoplasmic reticulum before being loaded onto MHC Class I^1^.

Which peptides are displayed by MHC Class I is dependent on binding preferences of hypervariable Class I genes (*HLA-A,-B,-C*)^2^. Far less is understood regarding how these peptides are generated from intracellular proteins by protein degradation pathways. It was initially reported that the bulk of MHC Class I peptide generation is dependent on ubiquitination and proteasomal degradation^3,4^. However, the necessity of ubiquitination^5,6^ and proteasomal degradation^7^ has been questioned, particularly in certain contexts such as ubiquitin-independent presentation of viral peptides^8^ and in individuals with MHC Class I alleles more likely to present proteasome-independent peptides^9^. Our goal here was to define the role of the ubiquitin-proteasome system in endogenous MHC Class I peptide presentation more conclusively at the level of individual peptides from proteins in the canonical proteome, although recent reports detail significant generation of peptides from the non-canonical proteome as well^10^. Our work, which represents the first application of quantitative mass spectrometry to estimate the contribution of the ubiquitin-proteasome system to MHC Class I peptide generation, will allow us to determine characteristics of proteins dependent on ubiquitin-proteasome system pathways for generation of MHC Class I peptides, as well as to determine whether specific components of these pathways^11^ play an outsized role in peptide generation.

Here, we present a systematic survey of the role of protein degradation pathways in MHC Class I peptide presentation. Technical optimizations were made to enable high-throughput and quantitative MHC Class I peptide mass spectrometry following chemical inhibition of protein degradation pathways, allowing us to identify peptides presented by MHC Class I that are dependent on these pathways. Most experiments were performed in immortalized B lymphoblasts, which abundantly present MHC Class I peptides while generally not cross-presenting peptides typically associated with MHC Class II (i.e., peptides from extracellular or endolyosomal pathway proteins^12^). Initially, we inhibited ubiquitination and proteasomal degradation, and observed that, in line with findings from initial reports^3,4^ but contrary to some more recent reports^5,6,7,9^, UPS pathways are largely required for MHC Class I peptide presentation. However, we also identified evidence for compensatory pathways that presented atypical peptides when these canonical pathways were inhibited, as well as a surprising number of proteasome-dependent substrates less dependent on ubiquitination. This prompted us to consider the relative contributions of autophagy as well as proteasome-associated factors such as p97/VCP to MHC Class I peptide generation. Our experiments demonstrate that these protein degradation pathways generate specific subsets of MHC Class I peptides, and also suggest that MHC Class I peptide display is relatively robust in the face of environmental insult to protein degradation pathways. Remarkably, partial inhibition of the proteasome also induces the presentation of a small number of atypical peptides. One of these atypical peptides identified by mass spectrometry was found to elicit an autologous immune response, suggesting that proteasome inhibition may have clinical utility in cancer immunotherapy contexts.

## RESULTS

### MHC Class I peptide mass spectrometry is improved by methodology for more accurate data normalization and removal of ‘background peptides’

We purified MHC Class I peptides as previously described^13^, and subsequently labeled samples on their N-termini with mass isomers to enable relative quantification by multiplexing in a single mass spectrometry run. Additionally, we developed a “spike-in standard” consisting of mouse cells expressing MHC Class I allele H-2K^b^ cross-presenting a defined peptide (SIINFEKL) and mixed this lysate at a 1:100 ratio based on cell number with our human cell lysates. This mouse MHC Class I•SIINFEKL standard was then co-purified with human MHC Class I complexes, by use of a H-2K^b^•SIINFEKL complex antibody mixed with a pan-human MHC Class I antibody at a 1:100 ratio (**Figure 1A**). The spike-in standard enables more accurate normalization to correct for potential sample loss during the peptide purification steps. Additionally, it allows for normalization when experimental treatment alters overall MHC Class I peptide presentation. This in particular is critical if one wishes to test a condition (e.g. proteasome inhibition) that has the potential to reduce presentation. To demonstrate the value of this normalization, B lymphoblasts were treated with the secretion inhibitor brefeldin A for 16h, which inhibits transport of MHC Class I to the cell surface^14^. MHC Class I peptides quantified by mass spectrometry were as expected in size **(Supplemental Figure 1A)**. Whereas brefeldin A reduced overall MHC Class I peptide presentation as evidenced by total peptide intensity measured by mass spectrometry, levels of the spike-in standard were not affected (**Figure 1B**). Statistical analysis of mass spectrometry data by widely used data normalization methods like total intensity normalization, however, obscured this treatment-induced decrease in total MHC Class I presentation (**Figure 1C**). Normalization to the spike-in standard avoids this artifact, while also correcting for any potential sample loss during processing steps.

**Figure 1.**
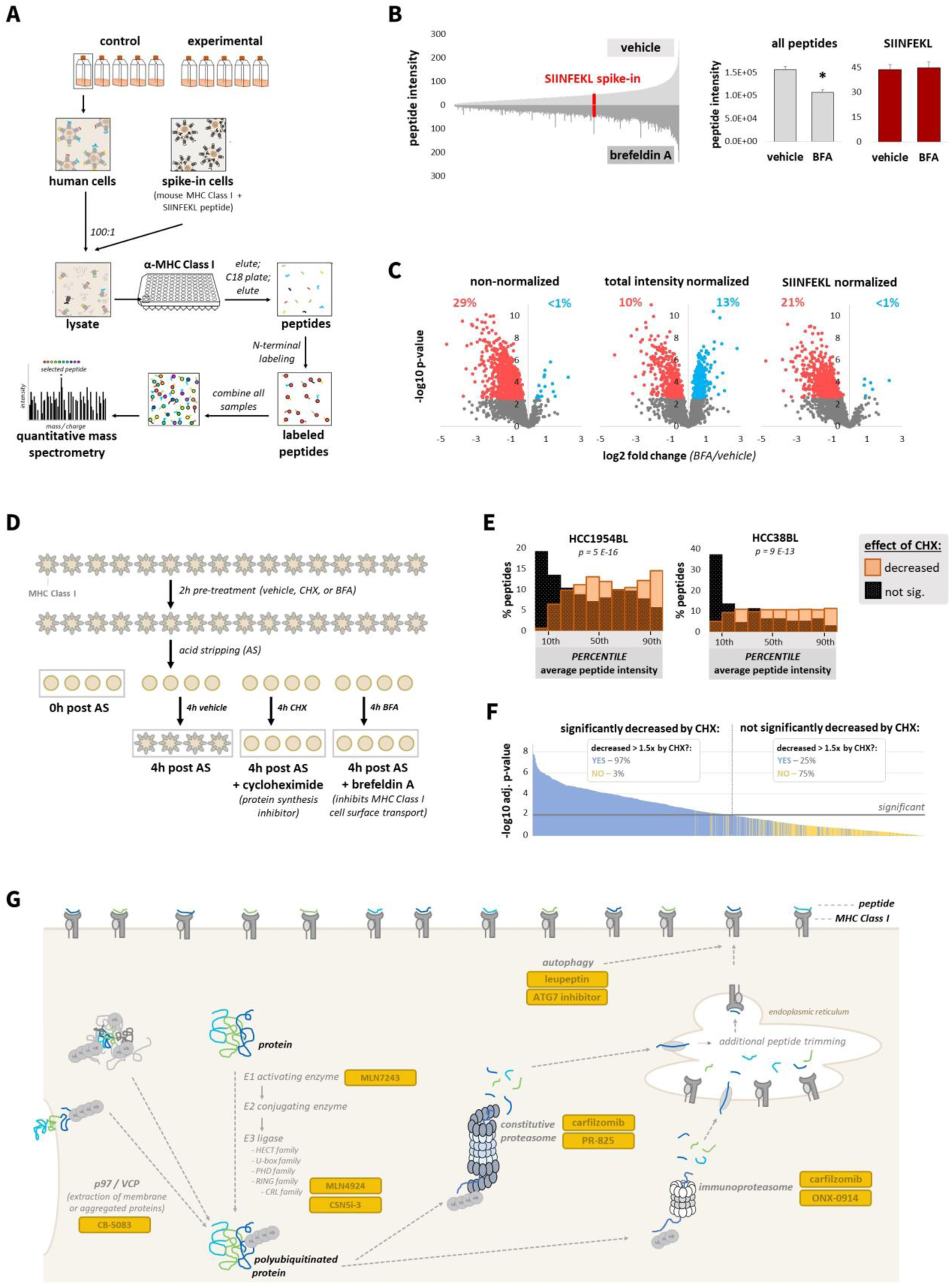
Improved methodology for MHC Class I peptide mass spectrometry. **(A)** Experimental design for mass spectrometry quantification of MHC Class I peptides. **(B)** Left: Average mass spectrometry individual peptide intensity from vehicle (upper) and brefeldin A (BFA) (lower) treated cells. SIINFEKL spike-in peptide is in red. Right: Summed total peptide intensity and SIINFEKL spike-in peptide intensity. * denotes p < 0.01 by t-test. **(C)** Changes in MHC Class I peptide presentation upon BFA treatment (red indicates significantly decreased; blue indicates significantly increased). Data are presented as non-normalized, total intensity normalized (each sample normalized to its summed peptide intensity), and normalized to the spike-in SIINFEKL peptide. Percent significant (increased and decreased) are marked above plots. **(D)** Experimental design to block MHC Class I peptide presentation; expected impact of treatments is depicted. **(E)** The average peptide intensity at 4h post acid stripping, representing constitutive antigen presentation, was determined for two groups: peptides significantly decreasing in response to CHX treatment, and those not significantly decreasing. Average peptide intensity percentiles, compared to all peptides identified, were determined and presented as histograms; p-value was determined by t-test. **(F)** The following groups were compared: peptides significantly decreasing in response to CHX treatment, and those not. Significance for each peptide is plotted. Another comparison using this dataset was performed, using just the first 2 replicates of the CHX treatment group. Peptides decreased greater than 1.5 fold in response to CHX in both replicates versus vehicle are plotted in blue, and all others plotted in yellow. In the legend, “decreased > 1.5 fold by CHX” means for both replicates. **(G)** Protein degradation pathways inhibited in our studies.

Ultimately, we aimed to perform experiments on cells treated with protein degradation pathway inhibitors for just a few hours due to the toxicity of inhibiting core pathways. Removal of pre-existing MHC Class I complexes by brief incubation of cells in mild acid solution (“acid stripping”)^15,16^ resulted in > 90% reduction in cell surface Class I complexes, with substantial recovery by 4h **(Supplemental Figure 1B)**. Cells were pre-treated for 2h with either brefeldin A, the translation inhibitor cycloheximide to block production of new proteins and MHC Class I complexes, or vehicle, and then acid stripped. Cells were immediately collected following acid stripping, or else cultured again for 4h with inhibitors (**Figure 1D**). We observed that total peptide intensity measured by mass spectrometry was markedly reduced following acid stripping or upon recovery in cycloheximide; this reduction was lesser upon recovery in brefeldin A **(Supplemental Figure 1C),** since loaded MHC Class I complexes can potentially be purified from the ER even if not transported to the cell surface^17^. Nevertheless, total peptide abundance was not as reduced as we expected even following acid stripping or cycloheximide treatment. We considered that this might be a limitation of mass spectrometry, such as background quantification. Additionally, the preference of mass spectrometry for quantifying abundant peptides^18^ likely biased towards selection of MHC Class I-bound peptides that decreased less in response to treatment. To better understand these “background peptides”, we specifically considered the effects of cycloheximide treatment, which was comparable to effects seen immediately upon acid stripping **(Supplemental Figure 1D)**. We observed that peptides that did not significantly decrease in response to cycloheximide were lower in intensity, leading us to suspect that many peptides near the limit of detection by mass spectrometry suffer from background quantification or are too variable to detect a significant decrease (**Figure 1E**). For most subsequent mass spectrometry experiments, we included 2 cycloheximide-treated replicates, and excluded from further analysis peptides not decreasing greater than 1.5 fold in response to cycloheximide in both replicates. This filter removed from further consideration a large number of “background peptides” that were not statistically significant (**Figure 1F**), without requiring the three or more replicates needed for a cutoff based on statistical significance. We then sought to apply this optimized methodology for quantitative MHC Class I peptide mass spectrometry to determine the role of diverse protein degradation pathways in MHC Class I peptide generation (**Figure 1G**).

### Inhibition of ubiquitination and proteasomal degradation results in a net decrease in MHC Class I peptide presentation, yet paradoxical increases in certain peptides

In four B lymphoblast cell lines with largely distinct MHC class I alleles **(Supplemental Figure 1E)**, we planned to inhibit ubiquitination with the ubiquitin activating enzyme E1 inhibitor MLN7243 and the proteasome with carfilzomib. We determined a dose of MLN7243 (500 nM) resulting in disappearance of most polyubiquitinated proteins with 4h of pretreatment **(Supplemental Figure 2A)**, and a dose of carfilzomib (1 µM) resulting in near complete proteasome inhibition with 1h of pretreatment **(Supplemental Figure 2B,C)**. Proteasomal degradation of known substrates was inhibited with this dose of carfilzomib **(Supplemental Figure 2D)**. We did not observe significant translational inhibition following MLN7243 or carfilzomib pretreatment **(Supplemental Figure 2E)**. For mass spectrometry experiments, cells were pretreated with these inhibitors, pre-existing MHC Class I peptides removed by acid stripping, and cells treated again with inhibitors for 4h. These treatments did not impact cell viability **(Supplemental Figure 2F)**. As measured by flow cytometry **(Supplemental Figure 2G)** and mass spectrometry **(Supplemental Figure 2H)**, inhibition of ubiquitination or proteasomal degradation reduced overall MHC Class I peptide presentation.

Reduction in MHC Class I peptide presentation upon inhibition of ubiquitination or proteasomal degradation was unlikely to be a nonspecific outcome of inhibitor toxicity, as we performed mass spectrometry on cells treated with similarly toxic or more toxic doses of the DNA damaging agent cisplatin **(Supplemental Figure 3A)**, and observed significant decreases in peptide presentation only at levels of cisplatin treatment substantially more toxic than that of proteasome inhibition **(Supplemental Figure 3B)**.

Whereas we observed an expected decrease in MHC Class I peptide presentation upon inhibition of ubiquitination or proteasomal degradation, a substantial fraction of peptides was not significantly decreased by these treatments (**Figure 2A**). Some of these peptides showed a trend towards decreasing that might become statistically significant if larger sample sizes were used. Nevertheless, we considered the possibility that MHC Class I peptides presented seemingly independent of the ubiquitin-proteasome system could be produced by alternative pathways. We determined the subcellular localization of peptide source proteins for MHC Class I peptides significantly decreased by both carfilzomib and MLN7243 treatment (“UPS-dependent”), and those not significantly decreased by either treatment (“UPS-independent”) (**Figure 2B**). Peptides not significantly decreasing were more likely to be derived from mitochondrial inner membrane proteins (**Figure 2C**), a region of the cell inaccessible to proteasomes, and were more likely to be longer in length than the typical 9mer **(Supplemental Figure 3C)**, suggesting that protein fragments produced by the proteasome are more optimal for MHC Class I peptide display.

**Figure 2.**
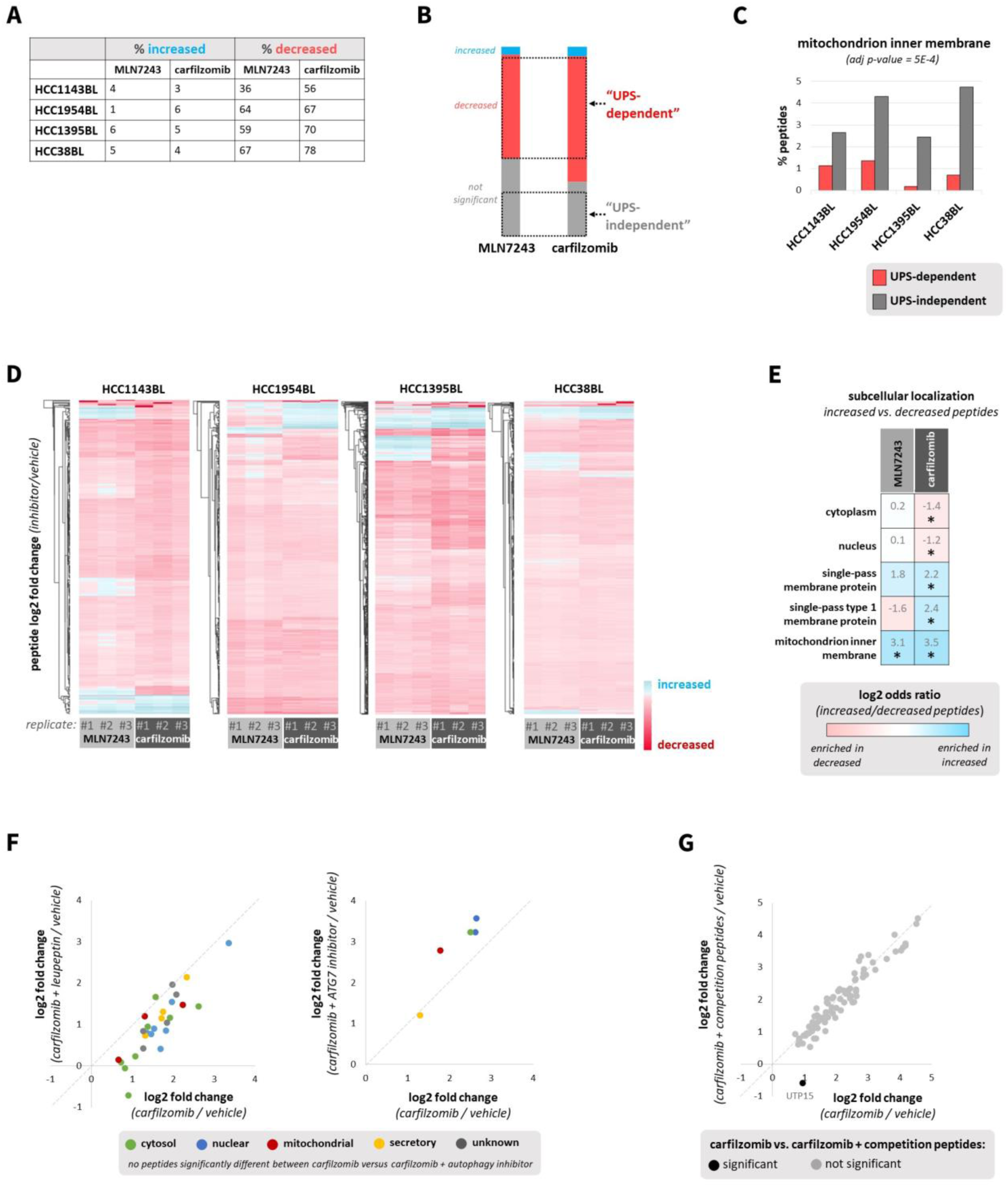
Inhibition of ubiquitination and proteasomal degradation decreases global MHC Class I peptide presentation, yet paradoxically increases certain peptides. **(A)** Percent MHC Class I peptides significantly increased or decreased by 4h MLN7243 (500 nM; 4h pretreatment) or carfilzomib (1 µM; 1h pretreatment) treatment versus vehicle. Peptides not decreased >1.5x by cycloheximide (25 µg/ml; 2h pretreatment) were excluded. **(B)** Criteria for classification of “ubiquitin-proteasome system (UPS)-dependent” and “UPS-independent” MHC Class I peptides. Bar charts represent combined data from cell lines in Figure 2A and are proportionally accurate. **(C)** Subcellular localization for UPS-independent and UPS-dependent peptides was obtained from UniProt, and a Cochran–Mantel–Haenszel test was used to test the enrichment of subcellular localization terms across all cell lines. **(D)** Heat map of peptides increased (blue) or decreased (red) by MLN7243 and carfilzomib treatment. Log2 fold change (inhibitor/vehicle) is depicted only for peptides significant in at least one treatment. **(E)** Enrichment of subcellular localization terms for peptides significantly increased versus peptides significantly decreased in cells treated with MLN7243 or carfilzomib. A Cochran–Mantel–Haenszel test determined the enrichment of subcellular localization terms across all cell lines; * indicates adjusted p-value < 0.05. **(F)** HCC1954BL cells were treated for 4h with carfilzomib (1 µM; 1h pretreatment) and/or leupeptin (50 µM; 1h pretreatment); similarly, they were treated with carfilzomib (1 µM; 1h pretreatment) and/or an ATG7 inhibitor (25 µM; 1h pretreatment). Only MHC Class I peptides significantly increased by carfilzomib are depicted. Scatterplots depict the log2 fold change of peptides for the following comparisons: carfilzomib versus vehicle (x-axis) and carfilzomib & leupeptin versus vehicle (y-axis), as well as carfilzomib versus vehicle (x-axis) and carfilzomib & ATG7 inhibitor versus vehicle (y-axis). **(G)** HCC38BL cells were treated for 4h with carfilzomib (1 µM; 1h pretreatment) and/or “competition peptides” known to bind HLA-A*03:01 and HLA-B*35:03 (20 ug/ml TIAPALVSK and YPTTTISYL, respectively; 1h pretreatment). Only MHC Class I peptides significantly increased by carfilzomib are depicted. Scatterplots depict the log2 fold change of peptides for the following comparison: carfilzomib versus vehicle (x-axis) and carfilzomib & competition peptides versus vehicle (y-axis).

Unexpectedly, we also observed that some peptides were increased by MLN7243 or carfilzomib treatment (**Figure 2A**). Significantly increased peptides often appeared to be inhibitor specific (**Figure 2D**), suggesting they were not produced by nonspecific cell stress pathways; in support of this, we also observed few increased peptides upon cisplatin treatment beyond rapidly induced proteins like HMOX1^19^ **(Supplemental Figure 3B).** By comparing significantly increased versus significantly decreased peptides across all cell lines, we observed that peptides significantly decreased in response to proteasome inhibition were more likely to come from cytosolic and nuclear proteins, whereas those that increased were more likely to be from single pass membrane proteins, particularly type 1 membrane proteins, and mitochondrial inner membrane proteins (**Figure 2E**). Peptides increased by inhibition of ubiquitination were also more likely to come from mitochondrial inner membrane proteins. Therefore, we suspected that MHC Class I peptides increased by proteasome inhibition, and perhaps inhibition of ubiquitination, were from proteins typically not exposed to the proteasome (e.g. mitochondrial and membrane proteins) and instead were perhaps degraded by extracytoplasmic proteases.

To test the hypothesis that MHC Class I peptides increased by proteasome inhibition were produced by autophagy or other forms of lysosomal degradation, we treated cells with carfilzomib in the presence or absence of the autophagy/lysosomal inhibitor leupeptin **(Supplemental Figure 3D)**, and quantified peptide presentation by mass spectrometry. Leupeptin treatment decreased overall MHC Class I peptide presentation, although less so than carfilzomib **(Supplemental Figure 3E)**, potentially due to its milder inhibition of select proteasomal activity^20^. Similarly, we also treated cells with carfilzomib in the presence or absence of an ATG7 inhibitor^21^ **(Supplemental Figure 3F)**, which decreased peptide presentation less significantly than leupeptin **(Supplemental Figure 3G)**. Most importantly, these autophagy inhibitors did not prevent the increased presentation of certain peptides observed with carfilzomib treatment (**Figure 2F**). Therefore, autophagy or lysosomal degradation, at least that able to be inhibited by the lysosomal enzyme inhibitor leupeptin or the ATG7 inhibitor blocking the induction of autophagy, does not appear to be the major source of MHC Class I peptides increased by carfilzomib treatment.

We appreciated that atypical MHC Class I peptides may still be able to be loaded onto free or recycled MHC Class I in the endolysosomal system, even if not produced by autophagy. Atypical antigen presentation pathways such as these may only become apparent in the absence of a larger supply of canonical MHC Class I peptides produced by the proteasome. To test this hypothesis, we treated a B lymphoblast cell line homozygous for HLA-A and HLA-B alleles with purified peptides identified previously by antigen mass spectrometry as able to bind these alleles (TIAPALVSK and YPTTTISYL for HLA-A*03:01 and HLA-B*35:03, respectively). Cells were also treated with carfilzomib, alone or in combination with these competition peptides. The competition peptides had predicted affinities for HLA-A and HLA-B higher than the majority of peptides identified, including the groups of all peptides identified by mass spectrometry and peptides significantly increased by carfilzomib treatment **(Supplemental Figure 3H)**. We confirmed that the competition peptides were loaded onto MHC Class I, and amongst the most abundant peptides detected by mass spectrometry. The competition peptides were also relatively abundant when compared with peptides significantly increased by carfilzomib treatment **(Supplemental Figure 3I)**. Therefore, we expect the competition peptides are sufficiently abundant and of high enough affinity to outcompete most peptides increased by carfilzomib if these peptides were loaded at the cell surface or in the endolysosomal system. As expected, treatment with the competition peptides did not substantially alter overall MHC Class I peptide presentation, suggesting they do not affect the presentation of canonical peptides produced by the proteasome and loaded in the endoplasmic reticulum **(Supplemental Figure 3J)**. Increased MHC Class I peptide presentation by carfilzomib was only significantly reduced in one instance by the competition peptides (**Figure 2G**), leading us to conclude that peptides induced by proteasome inhibition are also likely to be loaded in a conventional manner.

Therefore, we considered additional sources for the production of MHC Class I peptides increased by inhibition of ubiquitination or proteasomal degradation. Bioinformatic searches identified mitochondrial metalloprotease YME1L1 as enriched in being a putative binding partner of source proteins for the peptides that were increased by proteasome inhibition. **(Supplemental Figure 3K).** Cleavage by YME1L1 would allow for peptide sampling from mitochondrial inner membrane proteins that are protected from proteasomes. We also found that source proteins for peptides increased by inhibition of ubiquitination were more likely to be reported binders of ubiquitin-binding proteins HRS and GGA1 involved in protein sorting in the trans-Golgi network^22,23^ **(Supplemental Figure 3K).** It is possible some atypical sorting pathway directs these proteins producing MHC Class I peptides increased by MLN7243 to lysosomal (or other nonproteasomal) degradation. It is also possible that proteases in other cellular compartments (e.g. cytosolic, nuclear, endoplasmic reticulum) contribute to protein degradation and MHC Class I peptide generation in the absence of the ubiquitin-proteasome system.

### MHC Class I peptides only partially dependent on the ubiquitin-proteasome system share unifying characteristics

Given the differences in localization for MHC Class I peptides increased by inhibition of ubiquitination versus proteasomal degradation, we aimed to further define groups of peptides only partially dependent on the ubiquitin-proteasome system for their generation. We identified peptides that differed significantly in their response to MLN7243 or carfilzomib treatment in all cell lines, and grouped these peptides into “quadrants” **(Figure 3A; Supplemental Figure 4A,B).** These groupings can be generally understood as MHC Class I peptides particularly increased by proteasome inhibition (QI) or ubiquitination inhibition (QII), and peptides less ubiquitin-dependent (QIII) and those less proteasome-dependent (QIV). In support of these classifications, reported ubiquitin-independent proteasome substrate ODC1^24^ was observed in the expected QIII (**Figure 3A**). We then assessed whether subcellular localization and molecular function of MHC Class I peptide source proteins in these 4 quadrants were significantly different from peptides significantly decreased by both MLN7243 and carfilzomib. We observed that peptides in QI were more likely to come from single-pass type 1 membrane proteins, and peptide source proteins in QIV were more likely to be secreted proteins (**Figure 3B**). Peptides in QIII were more likely to be from transcription factors, some of which have been previously reported to be ubiquitin-independent substrates^25^. Peptides in QIV were more likely to be produced from MHC Class II proteins; the ubiquitin-dependent, proteasome-independent degradation of these proteins isn’t surprising, although the immunological significance of these MHC Class I peptides is unclear.

**Figure 3.**
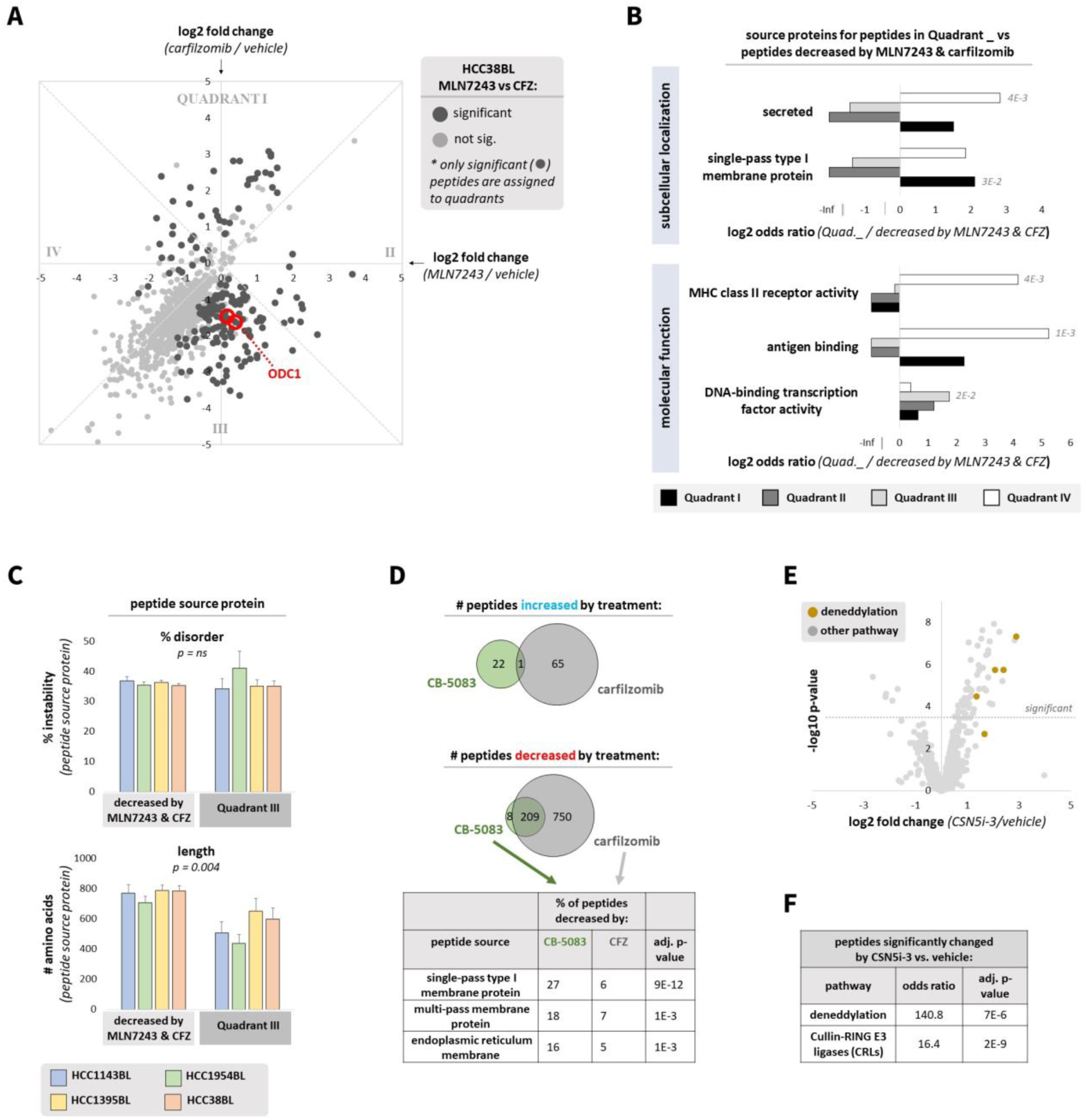
MHC Class I peptides only partially dependent on the ubiquitin-proteasome system share unifying characteristics. **(A)** HCC38BL cells were treated for 4h with MLN7243 (500 nM; 4h pretreatment), carfilzomib (1 µM; 1h pretreatment), or vehicle. Peptides not decreased >1.5x by cycloheximide (25 µg/ml; 2h pretreatment) were excluded. Scatterplot depicts the log2 fold change of peptides for the following comparisons: MLN7243 versus vehicle (x-axis), and carfilzomib versus vehicle (y-axis). Dashed perpendicular lines delineate “quadrants”; peptides from the reported proteasome-dependent, ubiquitin-independent substrate ODC1 are marked. **(B)** For each cell line, peptides significantly different between MLN7243 and carfilzomib treatment were assigned to quadrants as in Figure 3A. Peptides in each quadrant were compared to peptides significantly decreased by both MLN7243 and carfilzomib. A Cochran–Mantel–Haenszel test was used to test the enrichment of subcellular localization and molecular function terms from Uniprot across all cell lines; significant adjusted p-values are reported. **(C)** In each cell line, proteins assigned to QIII (significantly decreased more by carfilzomib versus MLN7243 treatment) were compared with proteins significantly decreased by both MLN7243 and carfilzomib treatment. The percent disorder and length of each source protein was calculated; significance was determined by Fisher’s method. **(D)** HCC1954BL cells were treated for 4h with CB-5083 (5 µM; 1h pretreatment), carfilzomib (1 µM; 1h pretreatment), or vehicle. Peptides not decreased >1.5x by cycloheximide (25 µg/ml; 2h pretreatment) were excluded. Venn diagrams depict the overlap of MHC Class I peptides significantly increased or decreased by CB-5083 and carfilzomib treatments. Significance of subcellular localization term enrichment (peptides decreased by CB-5083 versus peptides decreased by carfilzomib) was determined by Fisher’s exact test. **(E)** HCC1954BL cells were treated for 4h with MLN4924 (250 nM; 2h pretreatment), CSN5i-3 (1 µM; 2h pretreatment), or vehicle. Peptides not decreased >1.5x by cycloheximide (25 µg/ml; 2h pretreatment) were excluded. Volcano plot represents MHC Class I peptide presentation upon CSN5i-3 treatment versus vehicle. **(F)** Enrichment of deneddylation and Cullin-RING E3 ligases (CRLs) terms from Uniprot in peptides significantly changed by CSN5i-3 versus those not significantly changed was determined by Fisher’s exact test.

Proteasome-dependent, ubiquitin-independent proteins in QIII warranted additional investigation given their surprising number. Source proteins for MHC Class I peptides in QIII did not appear to be more disordered as a whole (**Figure 3C**) or more likely to be classified as intrinsically disordered **(Supplemental Figure 4C)**, which has been reported to target proteins to the proteasome independent of ubiquitination^26,27,28,29,30^. Peculiarly, the most striking feature of these proteins is that they are significantly shorter in length (**Figure 3C**). The finding of a large collection of MHC Class I peptides more dependent on proteasomal degradation than ubiquitination is suggestive of degradation by proteasomes lacking 19S regulatory particles (typically containing ubiquitin receptors and deubiquitinating enzymes), which does not directly require ubiquitination for protein degradation^31^.

Finally, we were interested in whether MHC Class I peptide generation generally requires the segregase/unfoldase p97/VCP. To test this, we treated cells with carfilzomib or the p97 inhibitor CB-5083 and assessed peptide presentation by mass spectrometry. As a whole, MHC Class I peptides appeared more dependent on proteasomal degradation than p97 for their generation (**Figure 3D**), with less than one fourth of proteasome-dependent peptides found to also be p97-dependent. However, p97-dependent peptides were preferentially enriched in being derived from multiple types of membrane proteins, as compared with proteasome-dependent peptides. This suggests the requirement of p97 in canonical MHC Class I antigen generation may be limited mostly to proteins requiring extraction from membranes, including endoplasmic reticulum membrane proteins as expected^32^, although cross-presentation of exogenous antigens by specialized cell types may also be p97-dependent^33^.

While inhibition of ubiquitination or proteasomal degradation produces atypical MHC Class I peptides (**Figure 2D, E**), sustained complete inhibition of these pathways across different tissue types is not clinically achievable. We considered whether partial inhibition of these pathways might be possible; for example, by inhibiting only the ubiquitination of certain proteins. Cullin-RING (CRL) E3 ligases are responsible for degradation of ∼20% of proteins targeted by the proteasome^34^ and can be selectively inhibited. We treated cells with the neddylation inhibitor MLN4924 and the COP9 signalosome inhibitor CSN5i-3, both of which alter the activity of CRL E3 ligases **(Supplemental Figure 4D).** Neither treatment had a large impact on MHC Class I peptide presentation **(Supplemental Figure 4E),** although it is possible that CRL substrates may be lower in abundance and thus less likely to be detected as peptides by mass spectrometry. Nevertheless, we did observe that CSN5i-3 treatment quite specifically increased presentation of CRL subunits and deneddylation components themselves as MHC Class I peptides (**Figure 3E,F**). This effect of CSN5i-3 towards increasing presentation of peptides from these pathways, which is not unexpected given that CSN5i-3 promotes turnover of select CRL complexes^35,36^, might have clinical applications in cancer patients with known mutations in these genes.

### Atypical MHC Class I peptide presentation can be elicited by partial proteasome inhibition

A method to elicit immunogenic MHC Class I peptide display in any patient, agnostic of mutation status, could have substantial clinical implications. We reasoned that this might be achievable via partial inhibition of the proteasome, if atypical peptides can be induced before dose-limiting toxicity is observed. Partial proteasome inhibition could also result in the production of novel peptides from proteins previously generating MHC Class I peptides but now with alternative sequence constraints, due to decreased activity of certain proteasome active sites (**Figure 4A**).

**Figure 4.**
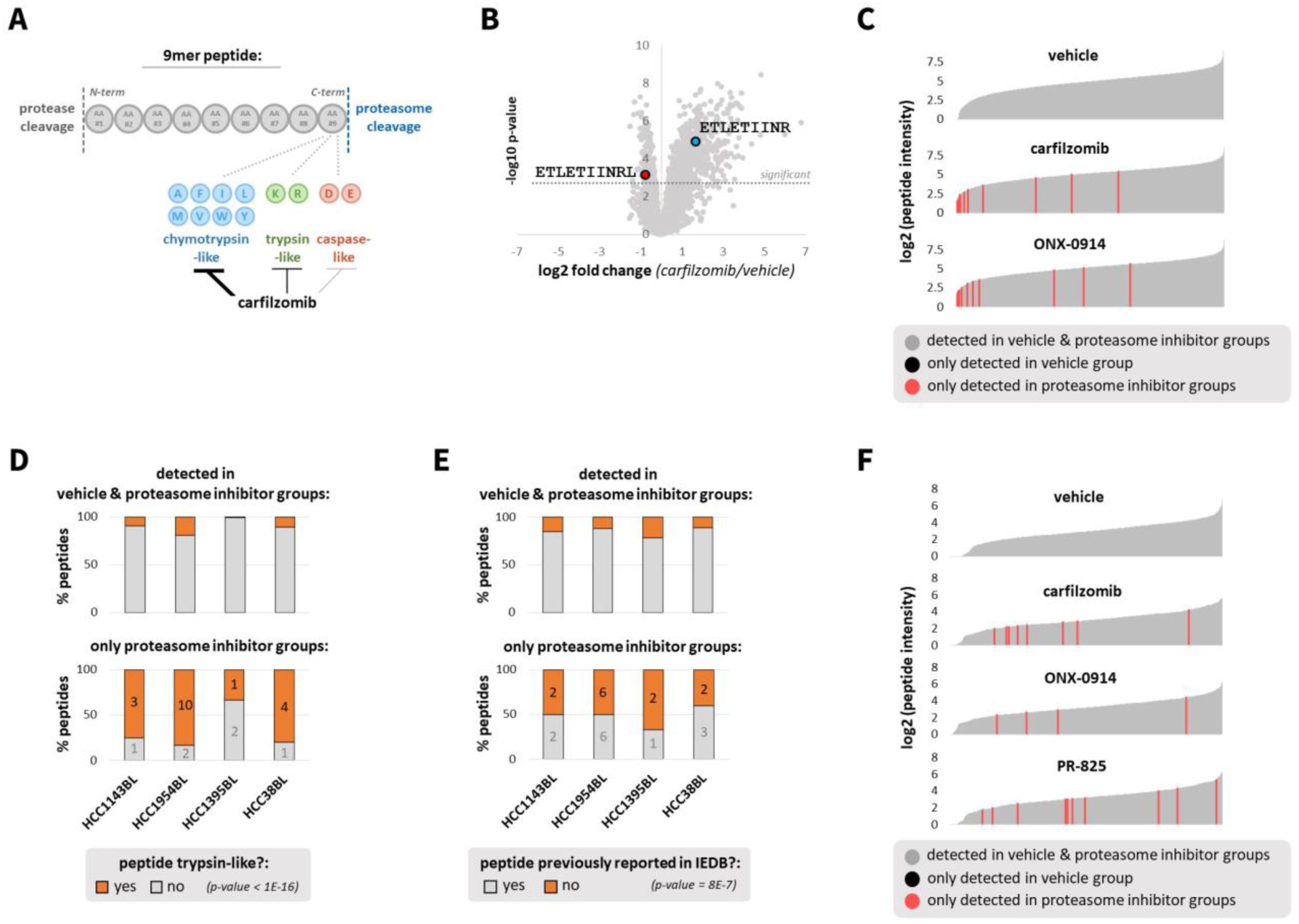
Atypical MHC Class I peptide presentation can be elicited by partial proteasome inhibition. **(A)** Impact of proteasome active site usage on MHC Class I peptide generation. A typical 9mer peptide is depicted, with the C-terminal amino acid assumed to be generated by proteasomal cleavage. Potential C-terminal amino acids generated by the 3 types of proteasome active sites (chymotrypsin-like, trypsin-like, and caspase-like) are indicated. Carfilzomib preferentially inhibits the chymotrypsin-like site. **(B)** HCC1954BL cells were treated with vehicle or carfilzomib (5 nM) for 48h to partially inhibit the proteasome, preferentially its chymotrypsin-like activity. Decreased production of a chymotrypsin-like peptide (ETLETIINRL) and increased production of an overlapping trypsin-like peptide (ETLETIINR) upon carfilzomib treatment is depicted. **(C)** HCC1954BL cells were treated for 48h with vehicle, or carfilzomib (5 nM) or immunoproteasome inhibitor ONX-0914 (25 nM) to partially inhibit the proteasome. Waterfall plots depict the log2 intensity of nonnormalized MHC Class I peptides detected in vehicle, carfilzomib, and ONX-0914 treated cells. Peptides not detected in vehicle treated cells (all 3 replicates) but detected in proteasome inhibitor treated cells (all 4 replicates) are marked in red. No peptides were detected in vehicle treated cells (all replicates) but not detected in either proteasome inhibitor treated cells (all replicates). **(D)** Cells were treated as described in Figure 4C. Plots show MHC Class I peptides detected in both vehicle and proteasome inhibitor treated cells (all replicates; top plot), and peptides detected only in proteasome inhibitor treated cells (all replicates of carfilzomib and ONX-0914 treatment; bottom plot). A Cochran–Mantel–Haenszel test was used to determine whether peptides only detected in proteasome inhibitor-treated cells were more likely to be “trypsin-like”. Numbers on bottom graph represent number of peptides. **(E)** Similar to Figure 4D, peptides bound to MHC Class I (HLA-A & HLA-B) previously reported in the Immune Epitope Database (IEDB) are shown in gray; those not previously reported are shown in orange. A Cochran–Mantel–Haenszel test was used to determine whether peptides only detected in proteasome inhibitor-treated cells were more likely not to have been previously reported in IEDB. **(F)** HCC1954 cells were treated with vehicle, carfilzomib (75 nM), ONX-0914 (200 nM), or constitutive proteasome inhibitor PR-825 (250 nM) for 48h. Waterfall plots depict the log2 intensity of nonnormalized MHC Class I peptides detected in vehicle, carfilzomib, ONX-0914, and PR-825 treated cells.

We were particularly interested in whether atypical MHC Class I peptides could be generated under conditions that mimic exposure to proteasome inhibitors in cancer therapy. For these experiments, we chose a dose of carfilzomib that inhibited the chymotrypsin-like site of the proteasome by approximately 75% **(Supplemental Figure 5A)**. This is comparable to the ≥75% inhibition of the chymotrypsin-like site observed in peripheral blood mononuclear cells of patients 1 h following carfilzomib treatment^37^. Because activity of the remaining proteasome active sites (trypsin-like and caspase-like) should remain largely intact under these conditions, we reasoned that overall protein degradation would be only modestly impaired. As a consequence, we anticipated that partial inhibition of the proteasome at the chymotrypsin-like site using clinically relevant concentrations of carfilzomib would alter the repertoire of peptides displayed by MHC Class I (**Figure 4A**).

We treated B lymphoblast cell lines for 48h with carfilzomib, or the immunoproteasome-specific chymotrypsin-like proteasome site inhibitor ONX-0914^38^, as immunoproteasomes are dominant in this cell line **(Supplemental Figure 5E).** Whereas complete proteasome inhibition greatly reduced MHC Class I peptide display **(Figure 2A; Supplemental Figure 2H)**, we observed that, with the exception of a small fraction of peptides that were significantly decreased **(Supplemental Figure 5C)**, that peptide display was generally unchanged or mildly upregulated by partial proteasome inhibition **(Supplemental Figure 5B).** As predicted, MHC Class I peptides significantly increased by partial proteasome inhibition targeting the chymotrypsin-like site, as compared to those decreased, had C-termini more likely to be produced by trypsin-like proteasomal cleavage **(Supplemental Figure 5D)**. This shifting from proteasomal chymotrypsin-like site activity to trypsin-like site activity could occasionally be observed for overlapping peptides (**Figure 4B**); we expect this would be observed more frequently in mass spectrometry experiments with deeper coverage.

While non-chymotrypsin-like peptides were more likely to be significantly increased by proteasome inhibition, most of these peptides were also detected in vehicle treated cells, and thus seem unlikely to be immunogenic. However, we were able to identify a small number of peptides only detected upon partial proteasome inhibition (**Figure 4C**). As expected, these peptides were more likely to be produced by the non-chymotrypsin-like sites of the proteasome (**Figure 4D**). They were also more likely not to have been previously reported as MHC Class I antigens in the Immune Epitope Database^39^ (**Figure 4E**), hence increasing their potential for immunogenicity.

The idea that MHC Class I peptides produced by mild proteasome inhibition, like that observed clinically, are immunogenic is difficult to reconcile with the fact that patients treated with these inhibitors do not appear to exhibit self-directed immune responses. However, proteasome inhibitors used clinically may also suppress the function of immune cells required for a response to atypical peptides. We considered that one way to bypass this limitation would be to treat solid tumor cells, which are constitutive proteasome dominant, with a proteasome inhibitor that specifically inhibits only the constitutive proteasome and thus would have little effect on immune cells that contain high levels of immunoproteasome.

To test this idea, we used HCC1954 breast cancer cells derived from the same patient as the HCC1954BL B lymphoblast cell line^40^. Unlike HCC1954BL, HCC1954 cells expressed approximately twice as many constitutive proteasome subunits as immunoproteasome subunits **(Supplemental Figure 5E,F)**. We aimed to treat these cells with carfilzomib, which targets constitutive and immunoproteasomes, as well as the immunoproteasome-specific inhibitor ONX-0914 and constitutive proteasome-specific inhibitor PR-825^41^ at doses that selectively inhibited the chymotrypsin-like site of the proteasome/immunoproteasome. As it is difficult to distinguish immunoproteasome versus constitutive proteasome inhibition *in vivo*, we assumed complete specificity of ONX-0914 and PR-825 and estimated the percent of chymotrypsin-like proteasome activity inhibition to be indicative of immunoproteasome/constitutive proteasome **(Supplemental Figure 5F)**. Complete inhibition of the chymotrypsin-like site with carfilzomib was too toxic to obtain; however, we expect to have inhibited the chymotrypsin-like activity of the immunoproteasome or constitutive proteasome completely with ONX-0914 or PR-825, respectively, with minimal off-target activity **(Supplemental Figure 5G).**

HCC1954 cells were treated with these doses of carfilzomib, ONX-0914, and PR-825 for 48h and MHC Class I peptide presentation assessed by mass spectrometry. As in HCC1954BL cells, overall peptide presentation was not reduced by these inhibitors **(Supplemental Figure 5H)**; instead, presentation of select peptides was enhanced **(Supplemental Figure 5I,J)**, particularly with carfilzomib and PR-825 treatment. Also as observed in HCC1954BL cells, certain peptides were only detected in proteasome inhibitor-treated cells (**Figure 4F**). That these peptides were observed in PR-825 treated cells further suggests that partial proteasome inhibition using constitutive proteasome-specific inhibitors may enhance immunogenic MHC Class I peptide presentation in solid tumor cells, while minimizing potential negative impacts on effector immune cells.

The potential therapeutic application of proteasome inhibition in cancer immunotherapy hinges upon the potential for novel MHC Class I-presented peptides to stimulate an immune response. To address this, we partially inhibited the proteasome in B lymphoblasts with carfilzomib or ONX-0914, and subsequently performed mass spectrometry to identify MHC Class I peptides that were either increased (“proteasome inhibitor-increased”) or exclusively detected (“proteasome inhibitor-specific”) in the presence of proteasome inhibitor (**Figure 5A**). Five peptides from each class **(Supplemental Figure 6A-D)** were chosen for further analysis in an *in vitro* peptide immunogenicity assay. In this assay, PBMCs from the same individual were stimulated with single purified peptides, and release of proinflammatory cytokines IFNγ and TNFα, as well as cytotoxic secretory granule proteins granzymes and perforin, were measured. We observed increased secretion of Granzyme A and Granzyme B in response to one proteasome inhibitor-specific peptide (**Figure 5B**).

**Figure 5.**
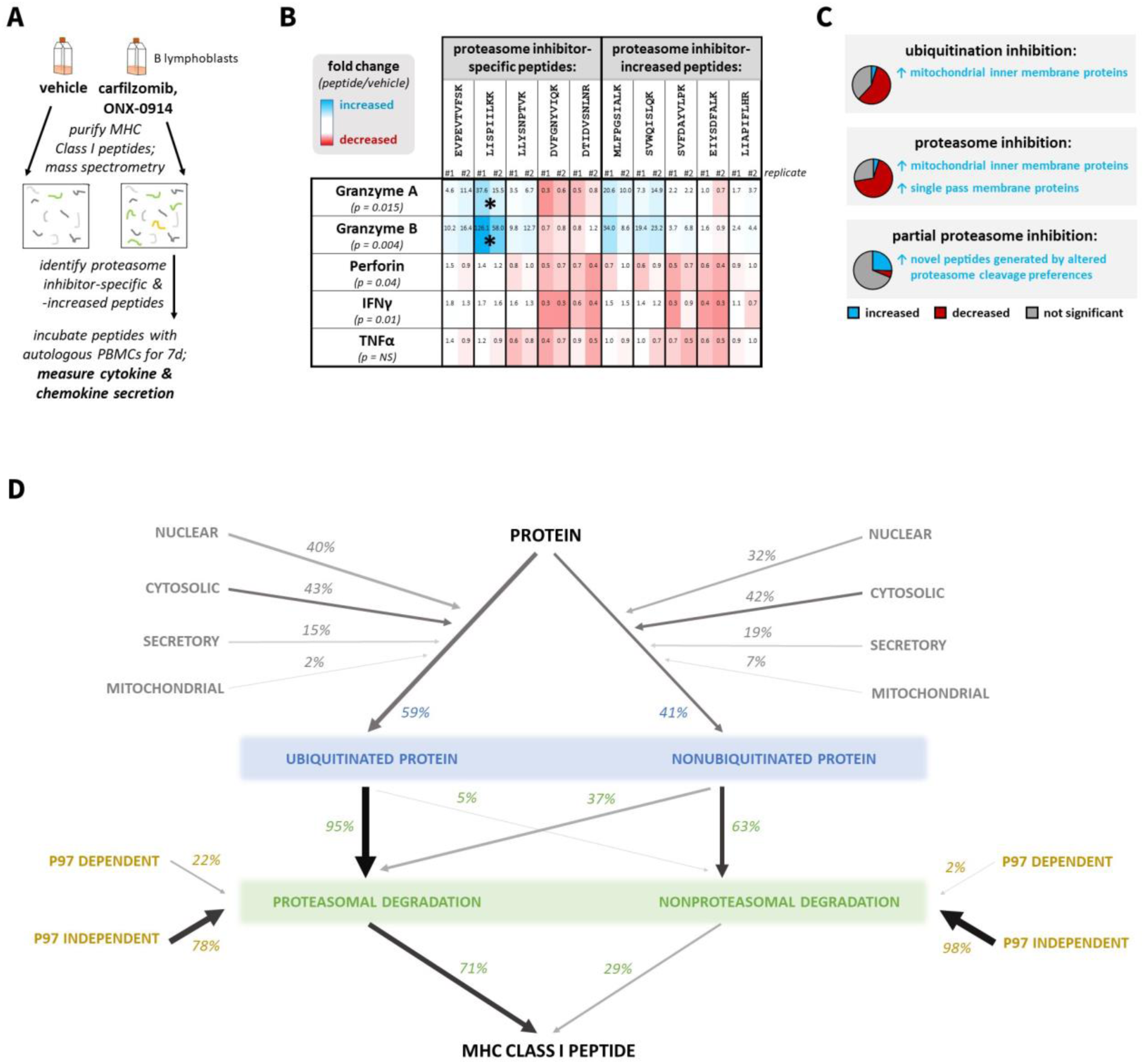
Diverse protein degradation pathways produce MHC Class I peptides, some of which demonstrate potentially immunogenicity when produced by compensatory pathways upon constitutive pathway impairment. **(A)** Experimental design for determining if proteasome inhibitor-induced or -specific peptides have potential immunogenicity. **(B)** Proteasome inhibitor-induced and - specific MHC Class I peptides were identified by mass spectrometry from B lymphoblasts treated for 48h with vehicle, or carfilzomib (5 nM) or immunoproteasome inhibitor ONX-0914 (25 nM) to partially inhibit the proteasome. Peripheral blood mononuclear cells (PBMCs) from the same individual were then incubated with these peptides for 7 d, and secretion of proinflammatory cytokines IFNγ and TNFα, as well as cytotoxic secretory granule proteins granzymes and perforin, measured. An ANOVA was performed to determine whether peptide treatment had a significant effect for each secreted factor; this p-value is listed (ns = not significant). When the ANOVA p-value was < 0.05, Dunnett’s test was performed to determine which peptide treatments were significant. * indicates p < 0.05. **(C)** Summary of impact of protein degradation pathway inhibition on MHC Class I peptide generation. Pie charts represent relative numbers of increased (blue), decreased (red), and not significantly changed (gray) peptides. Notable characteristics of significantly increased peptides (blue) are provided as text. **(D)** Quantitative summary of constitutive protein degradation pathways generating MHC Class I peptides. Numbers represent percent of peptides found to be significantly decreased in presentation by inhibition of indicated protein degradation pathways.

## DISCUSSION

Peptides presented on MHC Class I are selected from degradation products of endogenous proteins. These peptides play a key role in enabling the adaptive immune system to detect and eliminate cells that express ‘non-self’ proteins due to mutation or infection by bacteria or viruses. Here, we employed mass spectrometry to systematically quantify the contribution of the ubiquitin-proteasome system (UPS) to the repertoire of peptides presented by Class I molecules.

Although the UPS is widely thought to play a role in MHC Class I presentation, the nature and extent of its role has provoked some controversy. The strongest evidence for a role for the UPS comes from studies that quantify total levels of cell surface MHC Class I following chemical inhibition of the UPS, supplemented by analysis of presentation of select model MHC Class I-binding peptides in response to UPS inhibition^3–5^. While collectively these analyses indicate a role for the UPS in peptide presentation, they do not reveal the contribution of specific pathways within the UPS nor do they provide insight into the identities of the peptides that are dependent or independent of those pathways. Other groups have sought to address these points by using mass spectrometry to take a census of MHC Class I-bound peptides under different conditions. This has spawned unresolved ambiguity, particularly related to the role of the proteasome in MHC Class I peptide presentation. The first group investigating this question arrived at the surprising and unrefuted claim that there is little or no role for the proteasome in presentation of the majority of MHC Class I peptides^7^. Conversely, a later report from a second group supported a major role for the proteasome^42^, although they did not rationalize their results with those of Milner et al. and their claims were limited by the semi-quantitative nature of their mass spectrometry experiments.

To address this discrepancy, we first sought to address technical limitations of prior studies that quantified MHC Class I peptide presentation by mass spectrometry by developing new methodology to enhance the quality of our data (**Figure 1**), including the use of a spike-in MHC Class I peptide standard to enable more accurate normalization and a computational approach to exclude “background peptides” that are generally of low abundance and unresponsive to inhibitors that broadly block MHC Class I peptide presentation. Our approach enabled us to produce the most extensive catalog of UPS-dependent MHC Class I peptides generated to date. From our work across multiple cell lines, we concluded that most MHC Class I•peptide complexes require the proteasome (∼70%) and ubiquitination (∼60%) for their generation (**Figure 2**). An explanation for the discordant results regarding the role of the proteasome in MHC Class I peptide generation observed here versus previously reported^7^ is that the prior work prioritized minimizing proteasome inhibitor toxicity at the expense of achieving complete proteasome inhibition. Importantly, what we termed ‘partial proteasome inhibition’, for which we did not observe overall decreased MHC Class I peptide presentation (**Figure 4**), better reflects the extent of proteasome inhibition this group achieved.

One unexpected finding of our studies was that of the ∼70% of Class I-bound peptides that were dependent on proteasome activity, close to 20% were formed independently of ubiquitin conjugation activity (**Figure 3**). These peptides were enriched in being from transcription factors, which previous studies have suggested can serve as a source of ubiquitin-independent substrates^25^. Ubiquitination-independent peptides did not appear more likely to be from disordered proteins, but instead were derived from significantly shorter source proteins. The significance of this latter observation is unclear. It will be interesting to see if ubiquitin-independent peptides are generated by 26S proteasomes (as is the case for ubiquitin-independent ODC1^24^), free 20S proteasomes, which degrade nonubiquitinated hydrophobic peptides^43^ and oxidized or disordered proteins *in vitro*^30,44^ and *in vivo*^45^, or 20S proteasomes bearing alternative caps (e.g. PA28αβ, PA28γ, PA200)^46^.

Another component of the UPS that contributes to generation of a subset of Class I-bound peptides is the segregase/unfoldase p97/VCP. We found that p97 is required for the generation of ∼20% of presented peptides that are proteasome-dependent, particularly those from membrane proteins (**Figure 3**). This might be expected from the known roles of p97 in ER-associated degradation and membrane protein extraction^47^, although p97 may also be involved in cross-presentation of antigens from the endolysosomal system in specialized cell types. In contrast to p97, we observed little involvement of Cullin–RING ubiquitin ligase (CRL) activity in generation of MHC Class I•peptide complexes, despite these enzymes being involved in ∼20% of proteasome-dependent protein degradation^34^. Perhaps CRL substrates are, on average, too inabundant to be detected at the level of coverage achieved in our mass spectrometry experiments. Of note, CSN5i-3, a drug that binds and inhibits COP9 Signalosome (CSN) and thereby stimulates turnover of a subset of CRL subunits^35^, led to increased presentation of CSN and CRL subunits. Given that CRL subunits are relatively inabundant, this remarkable observation underscores the sensitivity and specificity of our methodology and its ability to detect more subtle perturbations to the UPS.

In addition to the prominent role for the UPS, a significant finding that emerged from our work was evidence for UPS-independent formation of MHC Class I•peptide complexes. We observed constitutive presentation of UPS-independent peptides disproportionately derived from mitochondrial inner membrane proteins, which are normally inaccessible to the UPS, although the mechanisms of their generation remain unclear. Another finding is that a smaller number of peptides (∼5%) are paradoxically increased by inhibition of UPS pathways. These peptides tend to be preferentially derived from mitochondrial proteins upon UPS inhibition, and single-pass transmembrane proteins upon proteasome inhibition alone. Although compensatory autophagy can occur in response to decreased proteasomal activity^48^, autophagy did not appear to be a major source of Class I-displayed peptides increased by proteasome inhibition, as induction of these peptides was not reduced by the lysosomal protease inhibitor leupeptin or an ATG7 inhibitor. We also determined that these peptides do not appear to be loaded in a compartment accessible from the cell exterior, arguing against a potential contribution from atypical antigen presentation pathways such as cross-presentation and MHC Class I recycling.

We then wondered whether increased presentation of atypical MHC Class I peptides upon proteasome inhibitor treatment could also occur in cancer therapeutic contexts where only partial proteasome inhibition is achievable^49,50^. To address this, we examined peptide presentation under conditions achieved upon carfilzomib treatment in blood cancer patients, in which the chymotrypsin-like site of the proteasome is inhibited by ∼75%, with much lesser inhibition of the trypsin-like and caspase-like sites^37^. Under these conditions, at least some protein degradation is expected to occur, but there should be a shift in the spectrum of peptides produced. Partial proteasome inhibition did not decrease overall peptide presentation; in fact, certain peptides were increased in presentation by partial proteasome inhibition, particularly those produced by non-inhibited proteasome active sites (**Figure 4**). We were especially interested in peptides detected only in proteasome inhibitor treated cells, which were rare (generally <10 detected per cell line) but significantly more likely not to have been previously reported. Of five peptides tested that were exclusively detected in proteasome-inhibited cells, one promoted immune system activation in assays using primary human immune cells (**Figure 5**). Additionally, we confirmed in a breast cancer cell line that proteasome inhibitor-specific peptides could be induced by proteasome inhibitors currently in the clinic (carfilzomib), as well as inhibitors selective for the immunoproteasome and constitutive proteasome (ONX-0914 and PR-825, respectively). While selective inhibitors of the constitutive proteasome have not been developed for clinical usage, our results suggest these could have application in cancer immunotherapy, since alternative and perhaps immunogenic peptide display could potentially be induced in solid tumor cells without impacting the viability or function of effector immune cells, which primarily express immunoproteasomes.

Notably, increased presentation of atypical MHC Class I peptides was also observed upon inhibition of additional UPS pathways. It is possible a larger number of unconventional compensatory protein degradation pathways, in the absence of canonical degradation pathways, could similarly generate potentially immunogenic MHC Class I peptides. For example, targeting a proteasomal substrate for endolysosomal degradation^51^ or a 26S proteasomal substrate for 20S proteasomal degradation^45^ is expected to alter the sequence of protein degradation fragments generated. It has also been reported that immunoproteasomes, which are inducible, generate MHC Class I peptides distinct from those generated by the constitutive proteasome^41,52^, although this is controversial^53^. In summary, we provide the most comprehensive quantitative analysis of the source of peptides loaded onto MHC Class I molecules. This work serves as a baseline resource for understanding the myriad mechanisms underlying the generation of the MHC Class I repertoire, which is fundamental to the ability of our adaptive immune systems to distinguish self from non-self. An interesting goal for future work will be to apply methods like those reported here to determine in greater mechanistic detail how individual pathways in the UPS contribute to the Class I peptide repertoire and to understand how genetic or environmental factors associated with the development of human inflammatory, infectious, or neoplastic diseases influence this repertoire.

## METHODS

### Study design

This study aimed to determine the contribution of diverse protein degradation pathways to generation of MHC Class I peptides using quantitative mass spectrometry. Multiplexed B lymphoblast samples treated with chemical inhibitors were generated for the following experiments: inhibition of MHC Class I secretion (HCC1954BL cell line; this cell line was used for all other experiments unless otherwise noted); inhibition of translation, MHC Class I secretion, and removal of cell surface MHC Class I by acidic peptide elution (HCC1954BL and HCC38BL cell lines); inhibition of ubiquitination and the proteasome (HCC1143BL, HCC1954BL, HCC1395BL, and HCC38BL cell lines); DNA damage and apoptosis induction; inhibition of autophagy in combination with proteasome inhibition (two separate experiments using leupeptin or an ATG7 inhibitor); inhibition of the proteasome in combination with competition peptide treatment (HCC38BL cells); inhibition of disaggregase p97/VCP; modulation of Cullin-RING E3 ligase activity; partial inhibition of the proteasome with carfilzomib and an immunoproteasome inhibitor (HCC1143BL, HCC1954BL, HCC1395BL, HCC38BL, and Cellero donor #287 cell lines). Multiplexed HCC1954 breast cancer cell line samples were also generated that were treated with carfilzomib, an immunoproteasome inhibitor, and a constitutive proteasome inhibitor for partial proteasome inhibition.

### Cell lines

HCC1143BL, HCC1954BL, HCC1395BL, HCC38BL, and HCC1954^40^ were purchased from ATCC and cultured in RPMI-1640 containing penicillin-streptomycin and 10% FBS. B lymphoblasts from donor #287 were purchased from Cellero and similarly cultured. For experiments, B lymphoblast cell lines were used at a concentration of approximately 1E6 cells/ml.

For certain experiments using B lymphoblasts, cells were pre-treated with protein degradation inhibitors or vehicle for times indicated, and cell surface MHC Class I removed by acid stripping. This was accomplished by bathing approximately 2E7 cells in 10 mL acid stripping buffer (0.132M citric acid, 0.06M sodium phosphate, pH 3.0) for 2 min on ice, followed by neutralization in 40 mL ice cold medium. Cells were washed once in medium prior to experimental treatment.

Cells were collected for mass spectrometry by centrifugation, which was preceded by trypsin detachment for adherent cell lines (HCC1954). Cell pellets were washed once in PBS and cell count and viability was assessed using a Vi-CELL XR.

### Inhibitors

Carfilzomib, MLN4924, and CB-5083 were purchased from Selleck Chemical. MLN7243 was purchased from Chemie Tek. Cycloheximide was purchased from Sigma-Aldrich. Brefeldin A and leupeptin were purchased from Alfa Aesar. Cisplatin was purchased from R&D Systems. CSN5i-3 and ONX-0914 were purchased from Fisher. PR-825 and the ATG7 inhibitor, compound #37^21^, were synthesized by WuXi AppTec.

### Spike-in SIINFEKL standard

Mouse DC2.4 dendritic cells were purchased from Millipore and maintained in RPMI-1640 containing 10% FBS, penicillin-streptomycin, & L-glutamine. Purified SIINFEKL peptide from ovalbumin protein (AnaSpec) was added to cells at 25 µg/ml for 4h. Cells were washed in PBS and stored as frozen pellets for future mass spectrometry experiments.

### Flow cytometry quantification of cell surface MHC Class I

Cells were washed twice in PBS. Approximately 1E5 cells were resuspended in 100 µl 2% FBS in PBS with anti-MHC Class I antibody (W6/32-PE) at a 1:50 dilution. Cells were incubated at 4°C for 30 min, then washed three times in 2% FBS in PBS. Flow cytometry was performed on a BD Symphony (MHC Class I expression following acid stripping) or a BD LSR II (MHC Class I expression following MLN7243, carfilzomib, and cycloheximide treatment in HCC1143BL and HCC1954BL cells). Experiments were gated on single cells, and 10,000 events captured per sample. Median fluorescence intensity of PE was calculated by BD FACSDiva.

### Proteasome activity assay

Approximately 5E5 cells were washed in PBS and lysed in 500 µl cold assay buffer [50 mM HEPES, pH 7.8, 10 mM NaCl, 1.5 mM MgCl_2_, 1 mM EDTA, 1 mM EGTA, 250 mM sucrose, 5 mM DTT]. Cells were vortexed and sonicated for 5s at 50% power, then centrifuged at 14,000 rpm for 10 min at 4°C. 50 µl lysate was loaded onto a 96 well plate on ice in triplicate for each proteasome activity probe. 200 µl assay buffer containing 2 mM ATP and 100 µM substrate (Suc-LLVY-AMC, Boc-LRR-AMC, Z-LLE-AMC, or Ac-nLPnLD-AMC) in DMSO was added to each well, and plates were incubated at 37°C for 1h. Plates were then imaged on a SpectraMax M5 fluorescent plate reader (excitation: 355 or 360 nM; emission: 460 nM). Fluorescence intensity was normalized to total protein content calculated by Bradford assay or total cell number.

### Quantification of translation

HCC1954BL cells were treated with 500 nM MLN7243 (4h), 1 µM carfilzomib (1h), 25 µg/ml cycloheximide (2h) or vehicle. For the final 30 min of treatment, cells were treated in methionine-deficient RPMI-1640 with 10% dialyzed FBS and penicillin-streptomycin. 1E6 cells were then labeled with 25 µCi/ml [^35^S]methionine (PerkinElmer NEG709A001MC) for 5 min. Unlabeled L-methionine (1 mg/mL) was added as previously described^54^ to terminate radiolabeling, and cells were centrifuged at 300×g for 1 min. Cells were washed with cold PBS containing 1 mg/ml L-methionine twice. Cells were then lysed in cold RIPA buffer for 5 min on ice and centrifuged at 14,000 rpm for 5 min. Proteins from supernatants were precipitated by adding 20% cold trichloroacetic acid on ice for 1h, followed by centrifugation at 14,000 rpm for 5 min. Pellets were washed with cold 5% trichloroacetic acid and subsequently cold acetone, with pelleting by centrifugation at 14,000 rpm for 5 min. Pellets were then resuspended in RIPA buffer and mixed with scintillation fluid. Incorporation of radiolabeled methionine was measured using a Beckman Coulter LS6500 scintillation counter set to read [^35^S] for 1 min.

### Purification of MHC Class I peptides

MHC Class I peptides were purified from frozen samples based on a protocol described previously^13^. Prior to purification, anti-human MHC Class I antibody (W6/32; Bio×Cell) was crosslinked to protein A-sepharose 4B beads by incubation for 1h at room temperature with shaking, followed by crosslinking in 20 mM dimethyl pimelimidate dihydrocholoride, 0.1 M sodium borate for 30 min at room temperature. Equal parts 0.2M ethanolamine, pH 8 was added and mixed for 5 min. Solution was removed from beads and beads were incubated with 0.2M ethanolamine, pH 8 for 2h at room temperature. Beads were washed three times in PBS and stored in an equivalent volume of PBS with 0.02% sodium azide. Similarly, antibody against mouse MHC Class I H-2K^b^ bound to SIINFEKL peptide (25-D1.16; Bio X Cell) was separately crosslinked to protein A-sepharose 4B beads. Anti-human MHC Class I antibody beads were mixed with anti-mouse MHC Class I•SIINFEKL beads at a 100:1 ratio.

Approximately 1E7 frozen human cells were lysed in 1 mL cold lysis buffer [PBS with 0.25% sodium deoxycholate, 0.2 mM iodoacetamide, 1 mM EDTA, protease inhibitor cocktail, 1 mM PMSF, and 1% octyl-beta-D-glucopyranoside]. Frozen mouse cells presenting the SIINFEKL spike-in standard were also lysed in this buffer. Cells were lysed on ice with occasional vortexing for 30 min, then lysates centrifuged at 14,000 rpm for 30 min at 4°C. During this time, a 96 well filter plate was washed with 200 µl acetonitrile, 200 µl 0.1% formic acid, and 2x with 200 µl of 0.1M Tris-HCl, pH 8. Plates were centrifuged at 200 rpm for 1 min at 4°C if needed.

For each experiment, cleared lysate volumes representing an identical number of cells were used. These lysates were mixed with mouse cells presenting SIINFEKL peptide at a ratio of 100:1 cells. 150 µl of antibody slurry was added to wells of the 96 well filter plate and washed with 200 µl lysis buffer. Lysates were then passed through wells containing antibodies by gravity flow. Wells were washed 4x with 200 µl cold 150 mM NaCl in 20 mM Tris-HCl, pH 8, 4x with 200 µl cold 400 mM NaCl in 20 mM Tris-HCl, pH 8, 4x with 200 µl cold 150 mM NaCl in 20 mM Tris-HCl pH 8, and 2x with 200 µl cold 20 mM Tris-HCl pH 8. Plates were centrifuged at 200 rpm for 1 min at 4°C to pass wash buffers through plate. During this time, a Waters Sep-Pak tC18 96 well plate was washed with 1 mL 80% acetonitrile in 0.1% formic acid, followed by 2 mL 0.1% formic acid. MHC Class I complexes were eluted from the antibody plate into the C18 plate with 500 µl 1% trifluoroacetic acid. The C18 plate was washed with 2 mL 0.1% formic acid, and MHC Class I peptides eluted with 500 µl 28% acetonitrile in 0.1% formic acid.

Purified peptides were dried using a GeneVac vacuum evaporator, and resuspended in 100 mM HEPES, pH 8. Peptides were N-terminally labeled using TMT labels (10 samples: TMT10plex; 11 samples: TMT10plex + TMT11-131C; 12-16 samples: TMTpro), and combined for a single mass spectrometry run. Peptides were dried and desalted using C18 10 µl ZipTips before analysis.

### LC-MS analysis using SPS-MS3

For most experiments, the entire sample was used for a single mass spectrometry run. Labelled peptides were subjected to LC-MS/MS analysis on an EASY 1000 nanoflow LC system coupled to a Fusion Tribrid Orbitrap mass spectrometer (Thermo Fisher Scientific) equipped with a Nanospray Flex ion source. Samples were directly loaded onto an Aurora 25cm x 75µm ID, 1.6µm C18 column (Ion Opticks) heated to 50°C. The peptides were separated with a 2-hour gradient at 350 nL/min as follows: 2–6% Solvent B (7.5 min), 6-25% B (82.5 min), 25-40% B (30 min), 40-98% B (1 min), and held at 98% B (15 min). Solvent A consisted of 97.8% H_2_O, 2% ACN, and 0.2% formic acid and solvent B consisted of 19.8% H_2_O, 80% ACN, and 0.2% formic acid. The Fusion was operated in data dependent mode. Spray voltage was set to 2.2 kV, S-lens RF level at 60, and heated capillary at 275°C. Full scan resolution was set to 120,000 at m/z 200 in Profile mode with an AGC target of 4 × 10^5^ and a maximum injection time of 50 ms. Precursor mass range was set to 400−1500 m/z and the isolation window was set to 0.7 m/z. For data dependent MS2 scans the cycle time was 3 sec, AGC target value was set at 5 × 10^4^, and intensity threshold was kept at 5 × 10^3^. CID fragmentation of precursors was performed with a fixed collision energy of 35%, activation time of 10 ms, and activation Q of 0.25. MS2 scans were then performed in the Orbitrap at 30,000 resolution in Centroid mode using auto scan range and a maximum injection time of 150ms.

Dynamic exclusion was enabled to exclude after 2 times for 60 sec with a mass tolerance of 10 ppm. A charge state filter was also applied to only include precursors of charge 2-5. Quantitative MS3 scans were then performed using Multi-notch Isolation. SPS precursors were selected from the mass range of 400-1600 m/z with a precursor ion exclusion window from -50 to +5 m/z. Quadrupole isolation of the precursor used an isolation window of 0.7 m/z while the ms2 isolation window was set to 3 m/z for up to 10 notches. The AGC target was 5×10^4^ and maximum injection time was set to 500 ms. HCD fragmentation was performed with fixed collision energy of 65% followed by Orbitrap detection at 50,000 resolution in Centroid mode using a scan range from 100-500 m/z.

### Mass spectrometry data search

Raw data were analyzed in Proteome Discoverer 2.4 (Thermo Scientific) using an unspecific (no-enzyme) search with the Byonic search algorithm (Protein Metrics) and UniProt human fasta file containing the spike-in peptide sequence SIINFEKL. PD-Byonic search parameters were as follows: precursor mass tolerance of 5 ppm, CID low energy fragmentation, fragment mass tolerance of 20 ppm, and a maximum of 2 common modifications and 1 rare modification. Cysteine carbamidomethylation and TMT-6 or TMTpro addition to peptide N-termini and lysine were set as static modifications. Methionine oxidation was a common dynamic modification (up to 2 per peptide) and deamidated asparagine or glutamine was set as a rare dynamic modification (only 1 per peptide). Precursor and charge assignments were computed from MS1. Byonic protein-level FDR was set at 0.01, while Percolator FDRs were set at 0.001 (strict) and 0.01 (relaxed). In the consensus workflow, peptide and PSM FDRs were also set at 0.001 (strict) and 0.01 (relaxed), with peptide confidence at least medium, lower confidence peptides excluded, minimum peptide length set at 7, remove peptides without a protein reference set to false, and apply strict parsimony set to true. Quantification was performed at the ms3 level using reporter ion S/N ratios with an average reporter S/N threshold of 35, a co-isolation threshold of 30%, and an SPS mass matches threshold of 70%.

### Mass spectrometry data statistical analysis

PSM output files from ProteomeDiscoverer were filtered for peptides not flagged in “Quan Info” as “ExcludedByMethod”. Only peptides 7-14 amino acids long were used. Peptides also needed to have an abundance quantitation for at least 3 replicates of 1 treatment to be used. After this filtering, missing values were rare; when they occurred, values were imputed from a Gaussian distribution centered on the bottom 1^st^ peptide intensity percentile with standard deviation of the median standard deviation of peptides in the bottom 10^th^ intensity percentile. PSMs for identical sequence peptides from the same UniProt ID, disregarding location of TMT labeling and modification, were averaged. Peptide intensities were then normalized to the spike-in standard SIINFEKL intensity.

A limma test was performed for statistical significance between treatments with R^55^. Results were considered significant when the adjusted p-value was < 0.01. After performing the limma test, negative values from Gaussian imputation were changed to the dataset minimum value, so that log2 fold changes could be calculated.

In certain experiments, an additional filter was applied: only peptides decreased at least 1.5 fold by cycloheximide treatment in both replicates. This filter was applied after the limma test was performed.

Heatmaps were generated using Seaborn’s *clustermap* function with no column clustering in Python.

### Peptide competition assay

MHC Class I peptides previously identified to bind HLA-A*03:01 (TIAPALVSK) and HLA-B*35:03 (YPTTTISYL) in HCC38BL cells were purchased as ≥95% pure synthetic peptides from Sigma-Aldrich. These peptides were solubilized in DMSO and used for cell treatments at 20 ug/ml concentration, in the presence of absence of carfilzomib. Predicted MHC Class I peptide affinity scores were determined by NetMHCpan EL 4.1 ran by the Immune Epitope Database (IEDB). Only HLA-A*03:01 and HLA-B*35:03 were included in the search. The peptide was assigned to the HLA allele with the highest score.

### Enrichment of subcellular localization and molecular function terms

The number of proteins associated with GO-Cellular Compartment and GO-Molecular Function terms was obtained from GeneTrail2^56^. Subcellular localization terms were also obtained from UniProt^57^. Simplified subcellular localization terms, excluding “unknown”, were obtained from SubCellBarCode^58^. Enrichment of terms for MHC Class I peptide nonredundant source proteins from one group of interest were compared with a second (e.g. nonredundant source proteins for peptides significantly increased by carfilzomib versus those significantly decreased). For experiments performed in a single cell line, enrichment of these terms was determined by Fisher’s exact test with Bonferroni multiple testing corrections. At least 5 proteins needed to be associated with the term in at least one of the groups of peptides to be considered. For experiments performed in multiple cell lines, enrichment of these terms across cell lines was determined by Cochran–Mantel–Haenszel test with Bonferroni multiple testing corrections. At least 5 proteins across all cell lines needed to be associated with the term in at least one of the groups of peptides to be considered.

### Enrichment of protein-protein interactions

Proteins reported to bind MHC Class I peptide source proteins were obtained from BioGRID^59^. Enrichment of interactors for peptide nonredundant source proteins from one group of interest were compared with a second (e.g. nonredundant source proteins for peptides significantly increased by carfilzomib versus those significantly decreased). Only low-throughput, physical interactors were used. Enrichment of these interactors across cell lines was determined by Cochran–Mantel–Haenszel test with Bonferroni multiple testing corrections. At least 5 proteins across all cell lines needed to be associated with the interactor in at least one of the groups of peptides to be considered.

### Assessment of protein degradation by cycloheximide chase

HCC1954BL cells were pretreated with vehicle or carfilzomib (1 μM) for 10 min. Cycloheximide (25 μg/ml) was then added, and cells collected at 0h, 1h, and 4h timepoints. Known ubiquitin-proteasome system substrates were obtained from UbiNet 2.0^60^. Of these proteins, the 5 proteins with the shortest reported half-lives in B cells^61^ were chosen for immunoblotting: JAK3, AMFR, JAK1, SQSTM1, ETV5, and IRF8. SQSTM1 was removed from consideration since autophagy can be induced by proteasome inhibition^48^. JAK3 did not decrease in response to cycloheximide in vehicle treated cells by the 4h timepoint and thus was not included.

### Immunoblotting

Cells were lysed in NP40 lysis buffer [50 mM Tris-HCl, pH 7.4; 150 mM NaCl; 1% NP40] for the following experiments: treatment of cells with leupeptin or the ATG7 inhibitor, treatment of cells with MLN4924 or CSN5i-3, and treatment of cells with vehicle or carfilzomib in combination with cycloheximide. Cells were lysed in lysis buffer used for MHC Class I peptide mass spectrometry [PBS with 0.25% sodium deoxycholate, 0.2 mM iodoacetamide, 1 mM EDTA, protease inhibitor cocktail, 1 mM PMSF, and 1% octyl-beta-D-glucopyranoside] for the following experiment: dose and time response treatment with MLN7243. Protein concentration was determined by BCA assay (MHC Class I peptide mass spectrometry lysis buffer) or Bradford assay (NP40 lysis buffer). Equivalent protein amounts were run on Tris-Glycine gels and transferred to nitrocellulose membranes using Teknova Electroblot Buffer. Membranes were blocked in 5% milk and primary antibodies used at 1:1000 concentration.

Antibodies used were to polyubiquitin (FK1; Enzo BML-PW8805-0500), K48 polyubiquitin (Millipore 05-1307), ubiquitinated H2B (Cell Signaling 5546S), β-actin (Cell Signaling 3700S), LC3B (Cell Signaling 2775S), p62 (Cell Signaling 5114S), CUL1 (Invitrogen 71-8700), AMFR (Cell Signaling 9590S), JAK1 (Cell Signaling 29261S), ETV5 (Cell Signaling 16274S), and IRF8 (Cell Signaling 83413T).

### Flow cytometry measurement of apoptosis

HCC1954BL cells were treated with vehicle, 50 µM cisplatin, or 500 µM cisplatin for 20h, acid stripped to remove pre-existing MHC Class I complexes, and treated again with vehicle, 1 µM carfilzomib, 50 µM cisplatin, or 500 µM cisplatin for 4h. Approximately 1E6 cells were stained with Annexin V-FITC and propidium iodide (PI), staining apoptotic and dead cells, respectively, using a Life Technologies Dead Cell Apoptosis kit. Cutoffs for Annexin V-FITC and PI staining were established using unstained cells. Cells were classified as “live” (below the cutoff for Annexin V-FITC and PI staining), “early apoptotic” (above the cutoff for Annexin V-FITC staining), “late apoptotic/necrotic” (above the cutoff for Annexin V-FITC and PI staining), and “other”. Experiments were gated on single cells, and 10,000 events captured per sample on a BD Symphony. Analysis was performed with FACSDiva.

### Fluorescent proteasome gels

1h prior to cell harvesting, cells were treated with 500 nM Me4BodipyFL-Ahx3Leu3VS. Cells were washed with PBS during harvesting and lysed in cold NP40 lysis buffer [50 mM Tris-HCl, pH 7.4, 150 mM NaCl, 1% NP40, & protease inhibitor cocktail]. Cells were lysed at 4°C for 30 min with rotation and centrifuged at 14,000 rpm for 5 min at 4°C. 10 µg of protein was loaded onto 16% Tricine SDS-PAGE gels and run for approximately 4-6h at 120 V with Tricine running buffer containing 1:400 NuPAGE antioxidant. Gels were imaged on a Typhoon FLA9500 fluorescent gel scanner (excitation: 473 nm; emission filter: BPB1 530DF20).

### Enrichment of deneddylation and Cullin-RING ligase terms after CSN5i-3 treatment

Proteins with deneddylation gene ontology terms were obtained from UniProt in November 2020. Proteins listed as Cullin-RING ligases (CRL) were obtained from UUCD^62^. Nonredundant source proteins for peptides significantly increased by CSN5i-3 treatment were compared with nonredundant source proteins for peptides not significantly changed by CSN5i-3 treatment. Enrichment of these terms in peptides significantly increased by CSN5i-3 treatment was determined by Fisher’s exact test with Bonferroni multiple testing corrections.

### Proteasome cleavage site prediction

The C-terminal amino acid was determined for MHC Class I peptides and classified as “chymotrypsin-like” (alanine, phenylalanine, isoleucine, leucine, methionine, valine, tryptophan, or tyrosine), “trypsin-like” (lysine or arginine), or “other” (all other amino acids)^63^. Enrichment of trypsin-like peptides in peptides increased by ONX-0914 versus those decreased was determined by Fisher’s exact test. Enrichment of trypsin-like peptides in peptides only detected after proteasome inhibitor treatment across all cell lines was determined by Cochran–Mantel–Haenszel test.

### Calculation of MHC Class I peptide source protein length and disorder

MHC Class I peptide source protein length was obtained from UniProt. The percent instability of peptide source proteins was obtained from Dryad^64,65^. Significance of differences in length and percent disorder between nonredundant proteins in QIII and those decreased by MLN7243 and carfilzomib treatment across all cell lines was determined by Fisher’s method combining t-test p-values from each cell line. Number of peptide source proteins considered to be intrinsically disordered was obtained from DisProt^66^. Significance of differences in fraction of nonredundant peptide source proteins characterized as intrinsically disordered in QIII versus those decreased by MLN7243 and carfilzomib treatment (“UPS-dependent”) across all cell lines was determined by Cochran–Mantel–Haenszel test.

### Peptide immunogenicity assay

Proteasome inhibitor-increased and proteasome inhibitor-specific MHC Class I peptides were identified by mass spectrometry as described above from B lymphoblasts (Cellero 1038-3217JY16 from donor #287) treated with 5 nM carfilzomib, 25 nM ONX-0914, or vehicle for 48h. The HLA-typing of this donor is as follows: HLA-A* 03:01/68:01, HLA-B*15:01/44:02, HLA-C*03:03/07:04. Peptides were identified that were significantly increased by both proteasome inhibitors (“proteasome inhibitor-increased”) or were only detected upon both carfilzomib and ONX-0914 treatment (“proteasome inhibitor-specific”). From each of these groups, 5 peptides were selected that were expected to bind either HLA-A*03:01 or HLA-A*68:01. Custom synthesized peptides were obtained from Thermo Scientific (1-4mg quantity, >95% purity, and no modifications).

All peptide samples and buffers were assessed for endotoxin levels by a LAL (Limulus amebocyte lysate) test with the Charles River Endosafe-PTS (Charles River) system prior to being used in the biological assay, as described previously^67,68^. The analysis was performed according to the manufacturer’s instructions. All samples were found to have endotoxin levels that were within the limit acceptable for these cell-based assays (< 1.50 EU/mL).

A PBMC assay evaluating cytokine/chemokine release in response to peptide stimulation was performed as previously described^67,68,69,70,71^ using human PBMCs (Cellero 5057MA21 from donor #287) from the same donor as EBV-immortalized B lymphoblasts used to identify proteasome inhibitor-increased and proteasome inhibitor-specific MHC Class I peptides by mass spectrometry. Cryopreserved PBMCs were thawed and plated on the day of the study. Viability and cell counts were assessed using a Vi-CELL XR. PBMCs were then plated at 3 × 10^5^ cells/well in a total volume of 200μL of RPMI growth media [RPMI-1640, 10% heat-inactivated fetal bovine serum, and 1% penicillin/streptomycin/L-glutamine (Life Technologies)] at 37°C in 96-well culture plates. Cells were then challenged in duplicate with MHC class I peptides at 10 ug/mL or water control of equivalent volume. Plates were incubated in a 5% CO2 incubator at 37°C for 7 days. Positive controls were cells treated with lipopolysacharide (Sigma Aldrich L4391), phytohemagglutinin (InvivoGen), or CD3-CD28 ImmunoCult (StemCell Technologies). Aliquots of cell culture supernatants were taken at 7 days post challenge for measurement of cytokine secretion using Luminex multiplex assays as described below. All cell derived supernatants were frozen and stored at -70°C prior to use.

Multiplex cytokine analysis on PBMC culture supernatants were performed using Milliplex multiplexed cytokine/chemokine panel kits (EMD Millipore) and a Luminex FlexMAP 3D instrument with the xPONENT version 4.0 software. The DropArray 96 well plate and LT Washing station from Curiox BioSystems was used to miniaturize all Luminex assays and protocols provided by Curiox BioSystems were followed. Cell culture supernatants were thawed and then centrifuged at 1200 rpm for 1 min before testing. Secretion of the following cytokines was quantified with the MILLIPLEX® Human CD8+ T Cell Magnetic Bead Panel Premixed 17 Plex - Immunology Multiplex Assay kit (EMD-Millipore): IL-6, TNF-α, MIP-1α, Granzyme B, IFN-γ, IL-5, Perforin, sFasL, IL-4, sCD137, IL-13, IL-2, Granzyme A, GM-CSF, sFas, IL-10, and MIP-1β.

The resulting raw data was further analyzed using Analyst software (VigeneTech, Inc.) to determine the cytokine concentration of the unknown samples against known concentrations of standards. Lipopolysacharide, phytohemagglutinin, and CD3-CD28 ImmunoCult were used as positive controls for the PBMC assay; all positive controls induced a strong response. The following cytokines and chemokines were chosen for further analysis: TNF-α, Granzyme B, IFN-γ, Perforin, and Granzyme A. An ANOVA was used to determine if peptide treatment had a significant effect. If the ANOVA p-value was < 0.05, Dunnett’s test was performed to determine which peptide treatments, if any, were significantly different from vehicle at α < 0.05.

### Statistics

A limma test was performed for statistical significance between treatments for MHC Class I peptide mass spectrometry experiments with R^55^. Results were considered significant when the adjusted p-value was < 0.01. For statistical comparison of multiple treatments with vehicle, an ANOVA was performed; if there was a treatment effect significant at p < 0.05, Dunnett’s test was performed. When t-tests were performed for multiple cell lines, Fisher’s method was used to determine a single metaanalysis p-value. To calculate enrichment of categorical terms in a single cell line, Fisher’s exact test with Bonferroni multiple testing corrections was performed. To calculate enrichment of categorical terms across multiple cell lines, a Cochran–Mantel–Haenszel test with Bonferroni multiple testing corrections was performed. Bar graphs with error bars represent average + SEM.

## Supplementary Materials

Table S1. Detailed description of MHC Class I peptide mass spectrometry datasets

Table S2. MHC Class I peptide presentation after brefeldin A treatment to block MHC Class I secretion in HCC1954BL cells

Table S3. MHC Class I peptide presentation after acid stripping, brefeldin A treatment, or translation inhibition in HCC1954BL and HCC38BL cells

Table S4. MHC Class I peptide presentation after ubiquitin, proteasome, or translation inhibition in HCC1143BL, HCC1954BL, HCC1395BL, and HCC38BL cells

Table S5. MHC Class I peptide presentation after apoptosis-inducing agent cisplatin treatment in HCC1954BL cells

Table S6. MHC Class I peptide presentation after proteasome, autophagy, or translation inhibition in HCC1954BL cells

Table S7. MHC Class I peptide presentation after proteasome inhibition, translation inhibition, or competition peptide treatment in HCC38BL cells

Table S8. MHC Class I peptide presentation after proteasome, p97, or translation inhibition in HCC1954BL cells

Table S9. MHC Class I peptide presentation after Cullin-RING E3 ligase modulation or translation inhibition in HCC1954BL cells

Table S10. MHC Class I peptide presentation after partial proteasome inhibition in HCC1143BL, HCC1954BL, HCC1395BL, and HCC38BL cells

Table S11. MHC Class I peptide presentation after partial proteasome inhibition in HCC1954 breast cancer cells

Table S12. MHC Class I peptide presentation after partial proteasome inhibition in Cellero donor #287 B lymphoblasts

## ACKNOWLEDGEMENTS

We thank Karl Beutner for insightful discussions on MHC Class I and Chris Spahr for help establishing mass spectrometry experiments. We thank Spiros Garbis and Tsui-Fen Chou from Caltech’s Proteome Exploration Laboratory for running mass spectrometry experiments. We thank Carlos Arbelaez and Mahshid Amini for help with peptide immunogenicity assays.

## Funding

J.L.M. was previously supported by a Life Sciences Research Foundation postdoctoral fellowship funded by Astellas Pharma and is currently supported by Amgen’s Postdoctoral Fellows Program. At the outset of this work R.J.D. was an Investigator of the Howard Hughes Medical Institute and was supported therefrom.

## Author contributions

J.L.M., J.A.J., J.R.L., R.V., and R.J.D. conceptualized the study. J.L.M., D.J.S., J.R.C., J.L., and S.P. performed experiments. J.R.C., M.K.J., and S.P. provided specialized expertise in immunology assays and B.L., A.M., and M.J.S. provided specialized expertise in mass spectrometry. J.L.M., D.J.S., J.R.C., J.A.J., M.K.J., J.R.L., B.L., M.J.S., B.V.L., R.V., and R.J.D. interpreted results. J.L.M. wrote the original manuscript. D.J.S., J.A.J., R.V., and R.J.D. revised the manuscript. R.J.D. directed the study.

## Competing interests

J.L.M., D.J.S., J.R.C., J.A.J., M.K.J., J.L., J.R.L., S.P., B.V.L., R.V., and R.J.D. are or were employees of Amgen, Inc., although this study was initiated while J.L.M. was a postdoctoral fellow in the lab of R.J.D. at California Institute of Technology. J.R.C holds patent for CAR T cell technology which is commercially licensed from City of Hope.

## Data and materials availability

Raw mass spectrometry and PSM analysis files from ProteomeDiscoverer are available from *(PRIDE link; not yet public)*. Processed mass spectrometry data is available in the Supplementary Materials.

---

**Supplementary Figure 1.**
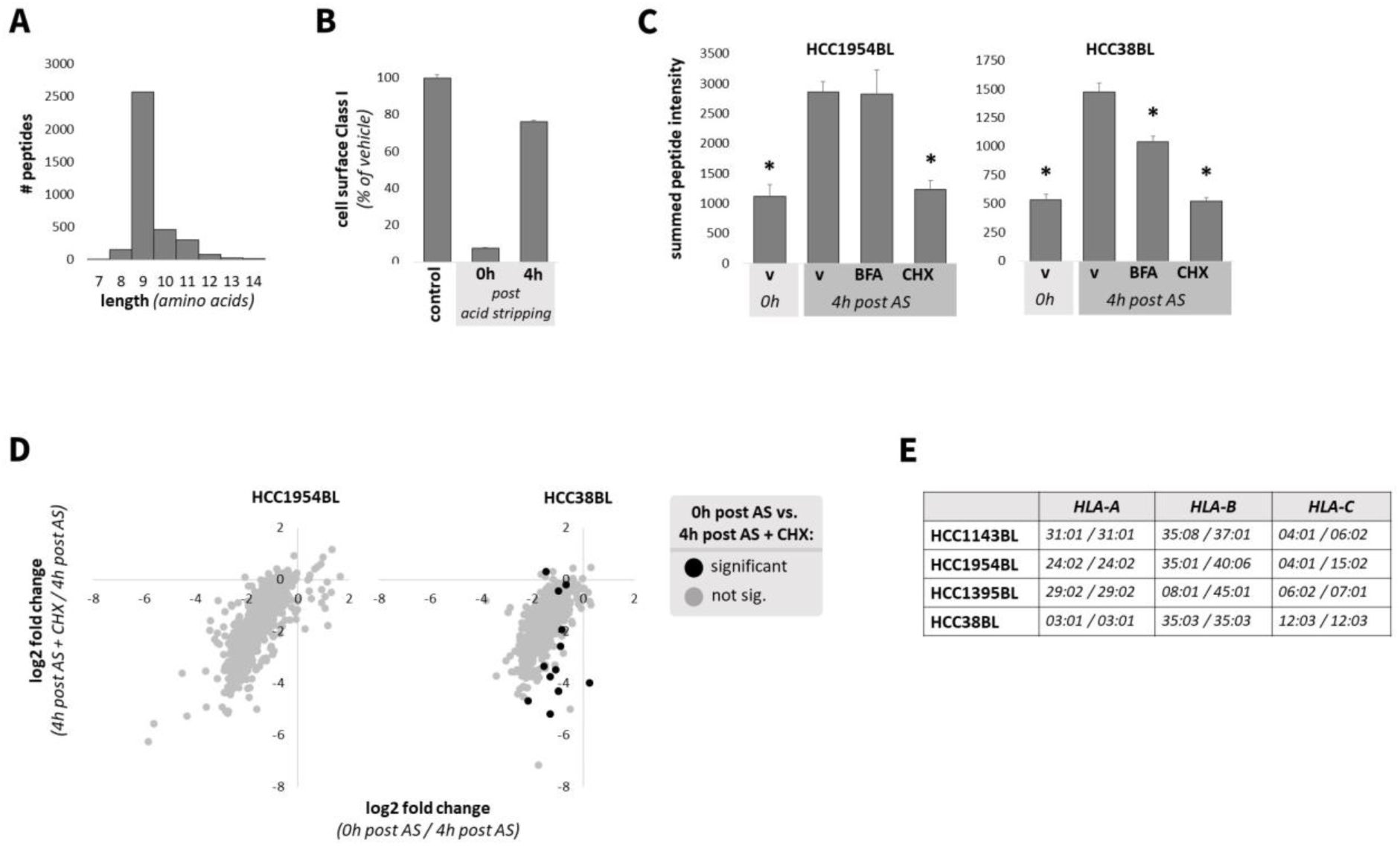
**(A)** Histogram of MHC Class I peptide length for all peptides identified in HCC1954BL cells treated with vehicle or 5 µM Brefeldin A for 16h. **(B)** Flow cytometry measurement of cell surface MHC Class I for untreated HCC1954BL cells (control), those collected immediately following mild acid elution to remove MHC Class I peptides (0h post acid stripping), and those collected 4h after acid stripping (n=3/group). **(C)** Cells were pre-treated with vehicle, cycloheximide (25µg/ml), or Brefeldin A (5 µM) for 2h, and pre-existing MHC Class I peptides removed by mild acid elution (“acid stripping”; AS). Cells were immediately collected after AS, or treated again with vehicle, cycloheximide, or BFA for 4h. Summed peptide intensity quantified by mass spectrometry is depicted. * indicates p < 0.01, as compared with 4h post AS + vehicle, by Dunnett’s test. **(D)** Scatterplots show the log2 fold change in displayed MHC Class I peptides from cells collected immediately after AS versus those collected 4h after AS (x-axis) or from cells collected 4h after AS with cycloheximide treatment versus those collected 4h after AS (y-axis). Black indicates peptides significantly different between cells collected immediately after AS versus those collected 4h after AS with cycloheximide treatment. **(E)** MHC Class I alleles for *HLA-A*, *-B*, & *-C* genes for B lymphoblast cell lines used.

**Supplementary Figure 2.**
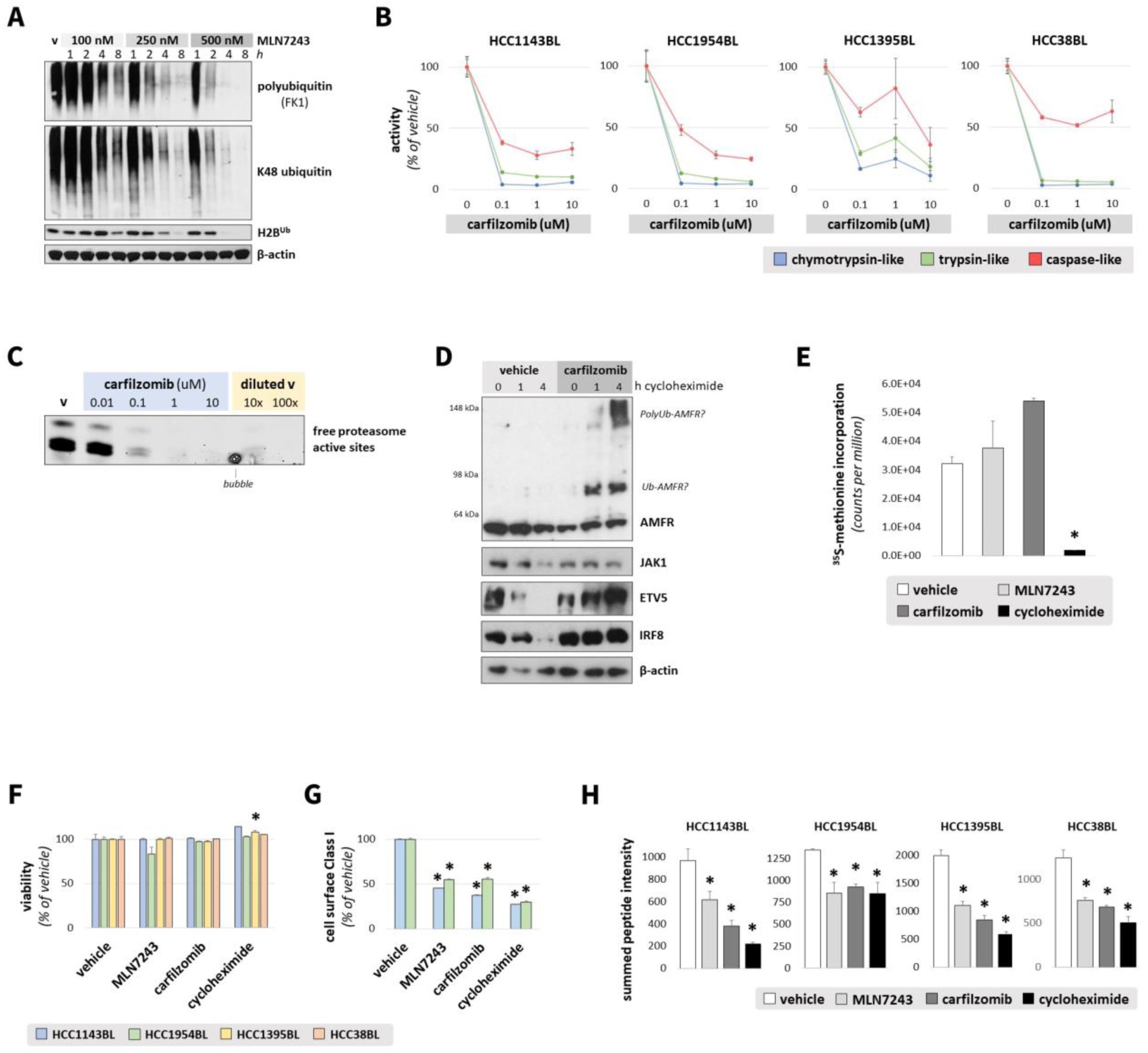
**(A)** HCC1954BL cells were treated with doses of the E1 ubiquitination inhibitor MLN7243 or vehicle (v) as indicated, for increasing lengths of time. Polyubiquitin, K48-linked ubiquitin, and ubiquitinated histone H2B (Lys120) were immunoblotted, with β-actin used as a loading control. **(B)** B lymphoblast cell lines were treated with doses of the proteasome inhibitor carfilzomib as indicated for 1h. Activity of the chymotrypsin-like, trypsin-like, and caspase-like sites of the proteasome was measured using fluorescent substrates (Suc-LLVY-AMC, Boc-LRR-AMC, and Ac-nLPnLD-AMC, respectively) incubated with cell lysates for 1h at 37°C. Percent inhibition was calculated relative to vehicle treated cells (n=3/group). **(C)** HCC1954BL cells were treated with vehicle (v) or increasing doses of carfilzomib for 1h. Free proteasome active sites were then labeled with 500 nM Me4BodipyFL-Ahx3Leu3VS for 1h. Cells were lysed and equivalent protein amounts (excluding rightmost two lanes) were loaded onto Tricine SDS-PAGE gels to visualize free proteasome subunits. Vehicle treated cells were also diluted 1:10 and 1:100 in lysis buffer, to represent 90% and 99% loss of free proteasome active sites, respectively. **(D)** HCC1954BL cells were treated with vehicle or 1 μM carfilzomib for 5 min. Cells were then treated with 25 μg/ml cycloheximide to inhibit translation and collected at timepoints listed. Equivalent protein amounts were loaded onto SDS-PAGE gels and immunoblotted for known ubiquitin-proteasome system substrates with reported short half lives in B cells. Β-actin was used as a loading control. **(E)** HCC1954BL cells were treated with vehicle, 500 nM MLN7243 (4h), 1 μM carfilzomib (1h), or 25 μg/ml cycloheximide (2h) (n=3/group). In the final 30 min of treatment, methionine-deficient media was used. Newly synthesized proteins were then labeled with [^35^S]methionine for 5 min, and incorporation of radiolabeled methionine measured in TCA precipitated proteins. * indicates p < 0.05, as compared with vehicle, by Dunnett’s test. **(F)** Viability of cells collected for mass spectrometry as measured by trypan blue staining, relative to viability of vehicle treated cells, for cells treated with MLN7243 (500 nM; 4h pretreatment), carfilzomib (1 µM; 1h pretreatment), and cycloheximide (25 µg/ml; 2h pretreatment) for 4h following acid stripping. * indicates p < 0.01 versus vehicle by Dunnett’s test. **(G)** Flow cytometry measurement of cell surface MHC Class I for HCC1954BL cells treated with MLN7243 (500 nM; 4h pretreatment), carfilzomib (1 µM; 1h pretreatment), and cycloheximide (25 µg/ml; 2h pretreatment) (n = 3/group) for 4h following acid stripping. * indicates p < 0.01 versus vehicle by Dunnett’s test. **(H)** Summed peptide intensity quantified by mass spectrometry for cells treated as described in Supplementary Figure 2F. * indicates p < 0.01 versus vehicle by Dunnett’s test.

**Supplementary Figure 3.**
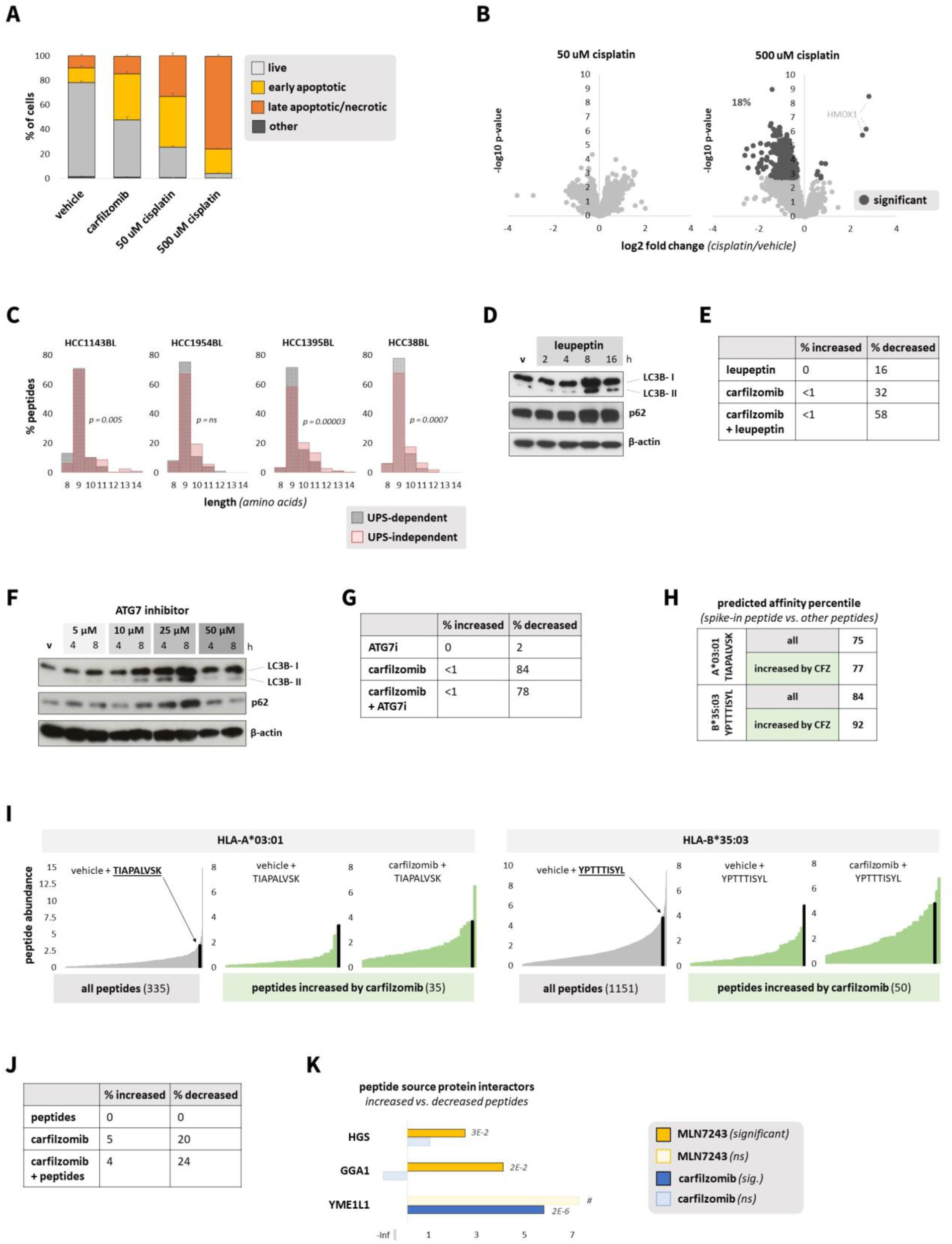
**(A)** Percent of HCC1954BL cells considered early apoptotic, late apoptotic/necrotic, live (not apoptotic or necrotic), or unclassifiable (“other”) after the following treatments for 4h post acid stripping: vehicle, 1 µM carfilzomib (1h pre-treatment), 50 µM cisplatin (20h pre-treatment), or 500 µM cisplatin (20h pre-treatment) (n=3/group). Staining for Annexin V was considered indicative of apoptosis and staining for propidium iodide indicative of necrosis. **(B)** Volcano plots representing quantitative changes in MHC Class I peptide presentation upon cisplatin treatment. Cells were treated for 4h post acid stripping with the following treatments: vehicle, 50 µM cisplatin (20h pre-treatment), and 500 µM cisplatin (20h pre-treatment). **(C)** Cells were treated with MLN7243 (500 nM; 4h pretreatment), carfilzomib (1 µM; 1h pretreatment), or cycloheximide (25 µg/ml; 2h pretreatment) for 4h following acid stripping. Peptides not decreasing more than 1.5 fold in response to cycloheximide were excluded. Histogram depicts peptide length for peptides not significantly decreased by MLN7243 and carfilzomib (“UPS-independent”) versus those significantly decreased by MLN7243 and carfilzomib (“UPS-dependent”). **(D)** HCC1954BL cells were treated with 50 µM leupeptin for timepoints indicated, and lysates were immunoblotted for LC3B, p62, and loading control β-actin. **(E)** HCC1954BL cells were treated for 4h with carfilzomib (1 µM; 1h pretreatment) and/or leupeptin (50 µM; 1h pretreatment); peptides not decreasing more than 1.5 fold in response to 25 µg/ml cycloheximide were excluded. Chart indicates percent of MHC Class I peptides significantly increased or decreased by each treatment. **(F)** HCC1954BL cells were treated with 5, 10, and 25 µM ATG7 inhibitor for timepoints indicated, and lysates were immunoblotted for LC3B, p62, and loading control β-actin. **(G)** HCC1954BL cells were treated for 4h with carfilzomib (1 µM; 1h pretreatment) and/or ATG7 inhibitor (25 µM; 1h pretreatment); peptides not decreasing more than 1.5 fold in response to 25 µg/ml cycloheximide were excluded. Chart indicates percent of MHC Class I peptides significantly increased or decreased by each treatment. **(H)** HCC38BL cells were treated for 4h with carfilzomib (1 µM; 1h pretreatment) and/or competition peptides known to bind HLA-A*03:01 and HLA-B*35:03 (20ug/ml TIAPALVSK and YPTTTISYL, respectively; 1h pretreatment). MHC Class I peptide affinity for HLA-A*03:01 and HLA-B*35:03 was predicted using NetMHCpan EL 4.1; the HLA allele with the highest predicted score was assigned to the peptide. The following groups of peptides were assessed, separated by predicted HLA allele: all peptides, and peptides significantly increased by carfilzomib treatment. The predicted affinity for the competition peptides (TIAPALVSK for HLA-A*03:01; YPTTTISYL for HLA-B*35:03) was similarly determined; chart indicates the percentile the predicted affinity of the competition peptide is in versus other peptides of the indicated group. Black bars represent the competition peptides. **(I)** Waterfall plots with gray bars depict the average MHC Class I peptide intensity for peptides after vehicle + competition peptide treatment, separated by predicted HLA allele binding. **(J)** Cells were treated as described in Supplementary Figure 3H. MHC Class I peptides not decreasing more than 1.5 fold in response to 25 µg/ml cycloheximide were excluded. Chart indicates percent of peptides significantly increased or decreased by each treatment. **(K)** Enrichment of proteins interacting with source proteins for MHC Class I peptides significantly increased versus peptides significantly decreased in cells treated with MLN7243 or carfilzomib. Cells were treated as in Supplementary Figure 3C. Protein interactors of source proteins for peptides significantly increased and peptides significantly decreased by MLN7243 and carfilzomib were obtained from BioGRID. A Cochran–Mantel–Haenszel test was used to test the enrichment of protein interactions across all cell lines; significant adjusted p-values are reported. “#” indicates insufficient interactions to calculate significance. ”-Inf” reflects no interactions in the increased peptide group.

**Supplementary Figure 4.**
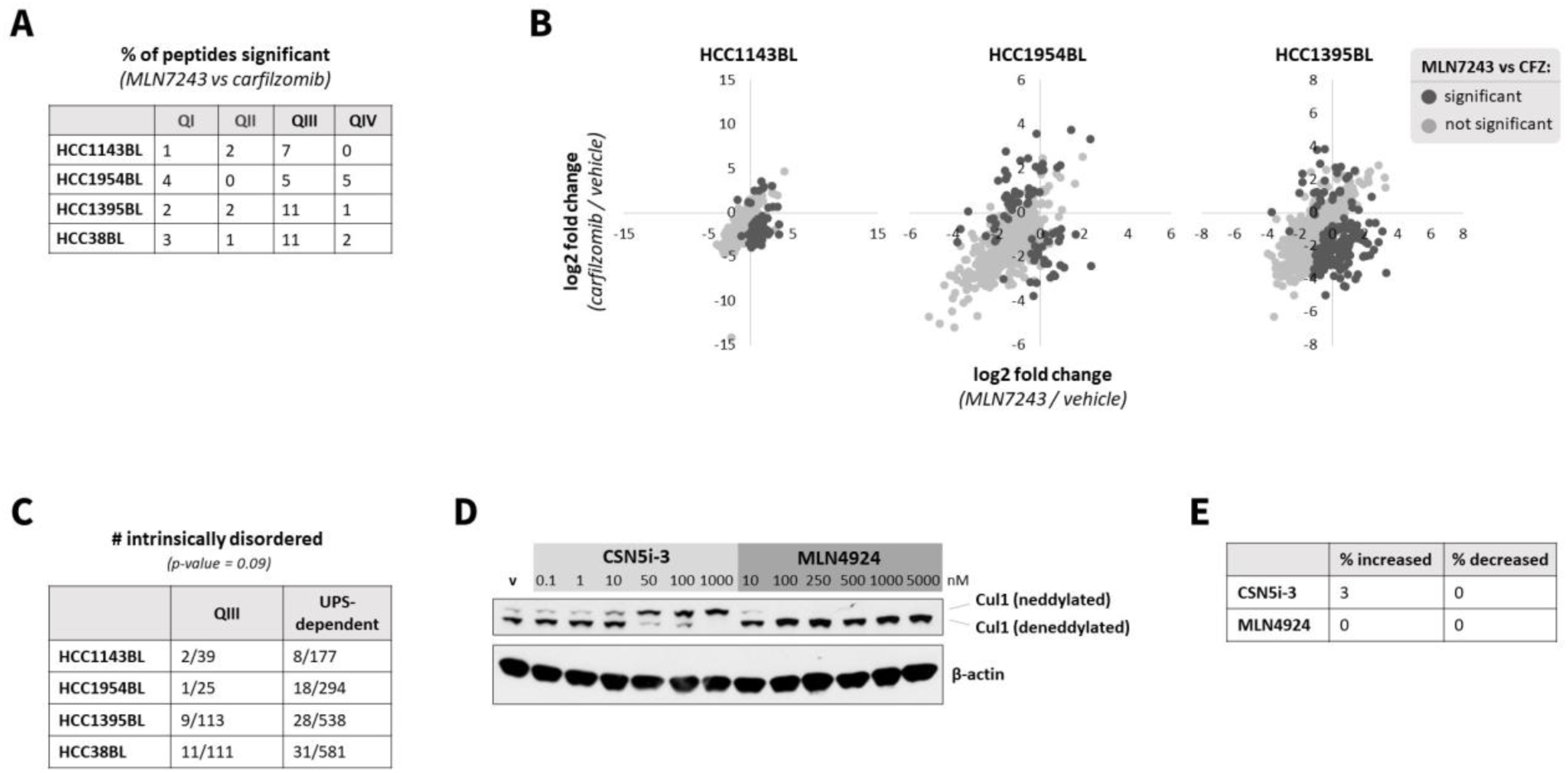
**(A)** Cells were treated for 4h with vehicle, MLN7243 (500 nM; 4h pretreatment), or carfilzomib (1 µM; 1h pretreatment). MHC Class I peptides not decreasing more than 1.5 fold in response to 25 µg/ml cycloheximide were excluded. Peptides significantly different between MLN7243 and carfilzomib treatment were assigned to “quadrants”, as depicted in Figure 3A. Table displays percent of peptides in each quadrant by cell line. **(B)** Cells were treated as in Supplementary Figure 3A. Scatterplot depicts the log2 fold change of MHC Class I peptides for the following comparisons: MLN7243 versus vehicle (x-axis), and carfilzomib versus vehicle (y-axis). **(C)** Number of proteins in QIII and “UPS-dependent” (significantly reduced by MLN7243 and CFZ) considered “intrinsically disordered” as classified by DisProt. A Cochran–Mantel–Haenszel test was used to determine whether the fraction of peptides considered intrinsically disordered across cell lines differed between the QIII and UPS-dependent groups. **(D)** HCC1954BL cells were treated with increasing doses of neddylation inhibitor MLN4924 or COP9 signalosome inhibitor CSN5i-3 for 2h. Lysates were immunoblotted for neddylated/deneddylated CUL1, with β-actin used as a loading control. **(E)** HCC1954BL cells were treated for 4h with vehicle, MLN4924 (250 nM; 2h pretreatment), or CSN5i-3 (1 µM; 2h pretreatment). MHC Class I peptides not decreasing more than 1.5 fold in response to 25 µg/ml cycloheximide were excluded. Chart represents percent of peptides significantly increasing and decreasing by MLN4924 and CSN5i-3 treatment.

**Supplementary Figure 5.**
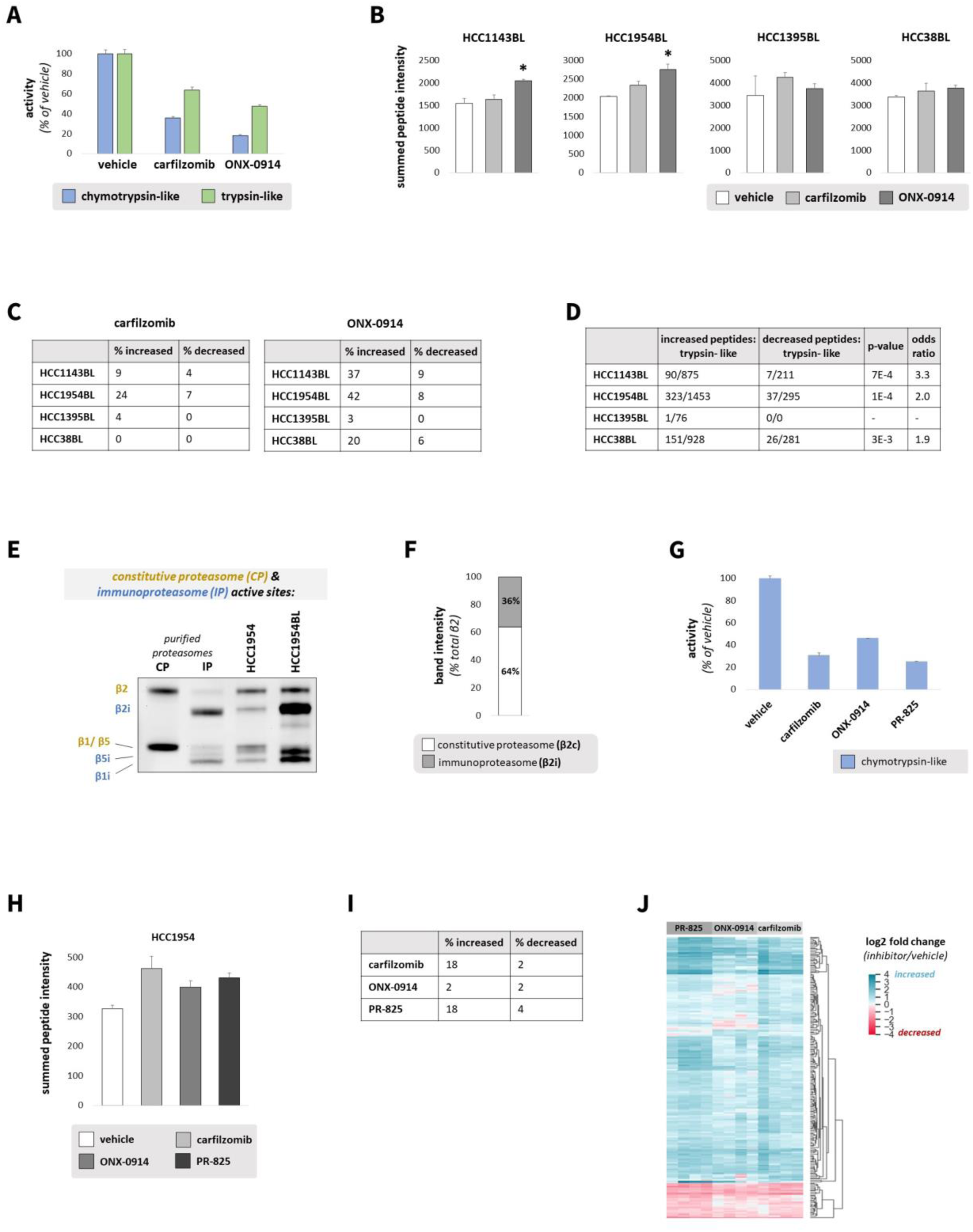
**(A)** HCC1954BL cells were treated for 48h with vehicle, carfilzomib (5 nM), or immunoproteasome inhibitor ONX-0914 (25 nM) to inhibit the chymotrypsin-like site of the proteasome. Activity of the chymotrypsin-like and trypsin-like sites of the proteasome was measured using fluorescent substrates (Suc-LLVY-AMC and Boc-LRR-AMC, respectively) incubated with cell lysates for 1h at 37°C. Percent inhibition was calculated relative to vehicle treated cells. **(B)** Cells were treated with vehicle, carfilzomib (5 nM), or ONX-0914 (25 nM) for 48h. Graph depicts summed MHC Class I peptide intensity; * indicates p < 0.01 by Dunnett’s test. **(C)** Percent of MHC Class I peptides significantly increased or decreased by carfilzomib or ONX-0914 treatment across cell lines. **(D)** The number of MHC Class I peptides considered “trypsin-like” (containing a C-terminal lysine or arginine) was determined for peptides significantly increased and decreased by ONX-0914 treatment versus vehicle. Significance was determined by Fisher’s exact test. **(E)** HCC1954 and HCC1954BL cells were treated with 500 nM Me4BodipyFL-Ahx3Leu3VS for 1h to label proteasome active sites. Equivalent protein amounts were loaded onto Tricine SDS-PAGE gels to resolve constitutive proteasome (yellow) and immunoproteasome (blue) subunits. Purified constitutive proteasome (CP) and immunoproteasome (IP) were also analyzed. **(F)** Quantification of proteasome active site fluorescent gel band intensities as a measure of immunoproteasome to constitutive proteasome ratios in HCC1954 cells. Intensities of β2 (constitutive proteasome trypsin-like site) and β2i (immunoproteasome trypsin-like site) bands were calculated from the gel in Supplementary Figure 5E. Intensity ratios are depicted. **(G)** HCC1954 cells were treated with vehicle, 75 nM carfilzomib, 200 nM ONX-0914, or 250 nM constitutive proteasome inhibitor PR-825 for 48h. Activity of the chymotrypsin-like site of the proteasome was measured using a fluorescent substrate (Suc-LLVY-AMC) incubated with cell lysates for 1h at 37°C. Percent inhibition was calculated relative to vehicle treated cells (n=3/group). **(H)** HCC1954 cells were treated with vehicle, carfilzomib (75 nM), ONX-0914 (200 nM), or PR-825 (250 nM) for 48h. Graph depicts summed MHC Class I peptide intensity. **(I)** Percent MHC Class I antigens significantly increased or decreased by carfilzomib, ONX-0914, or PR-825 in HCC1954 cells is listed. **(J)** Heat map shows log2 fold change (proteasome inhibitor/vehicle) for MHC Class I peptides in HCC1954 cells for peptides significant in at least one treatment.

**Supplementary Figure 6.**
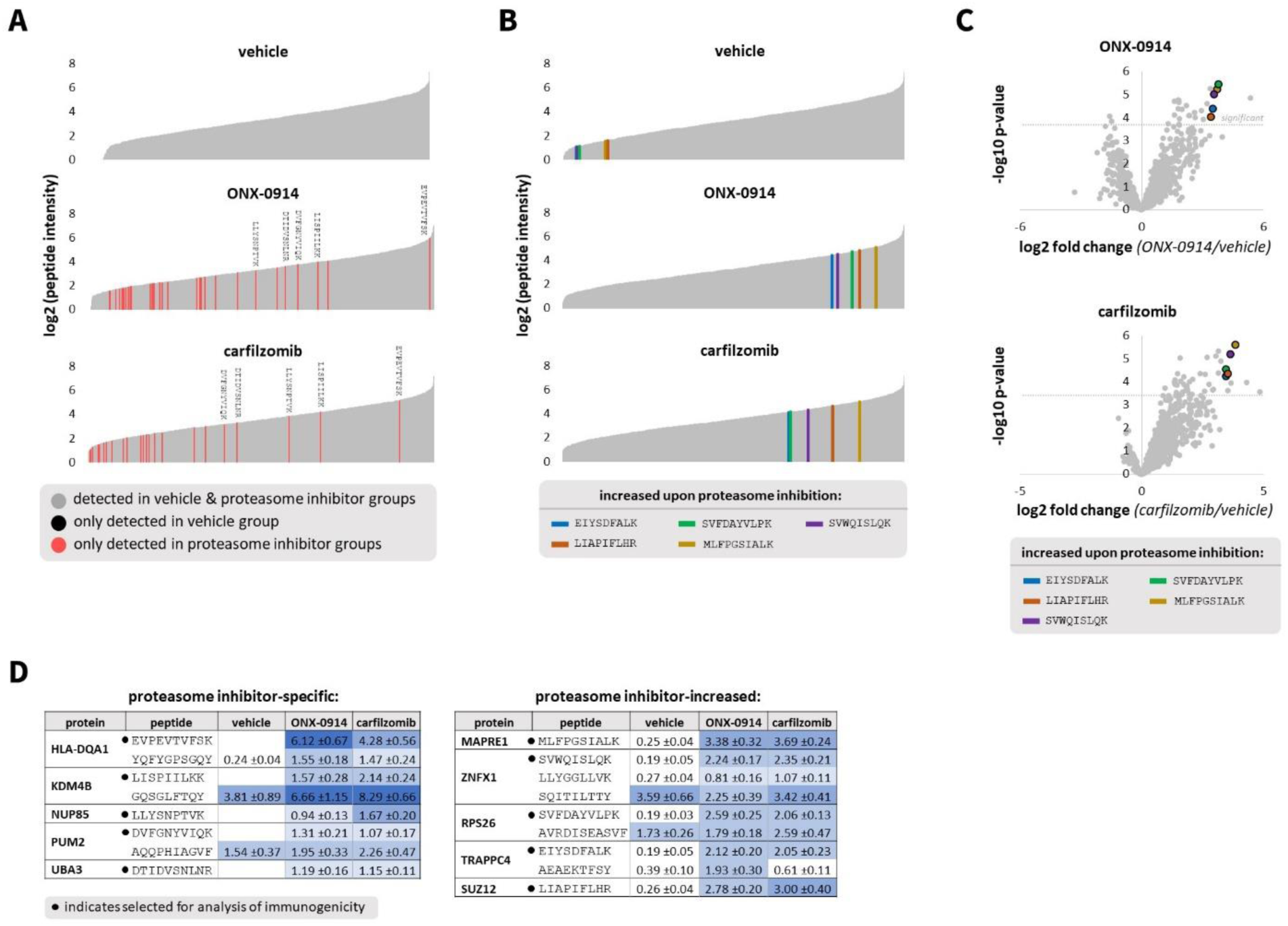
**(A)** B lymphoblasts (Cellero donor #287) were treated for 48h with vehicle, or carfilzomib (5 nM) or immunoproteasome inhibitor ONX-0914 (25 nM) to partially inhibit the proteasome. Waterfall plots depict the log2 intensity of nonnormalized MHC Class I peptides detected in vehicle, carfilzomib, and ONX-0914 treated cells. Peptides not detected in vehicle treated cells (all 3 replicates) but detected in proteasome inhibitor treated cells (all 4 replicates) are marked in red. No peptides were detected in vehicle treated cells (all replicates) but not detected in either proteasome inhibitor treated cells (all replicates). Peptide sequences listed were chosen for analysis of immunogenicity. **(B)** Waterfall plots depict the log2 intensity of nonnormalized MHC Class I peptides detected in vehicle, carfilzomib, and ONX-0914 treated B lymphoblasts (Cellero donor #287). Five peptides significantly increased by carfilzomib and ONX-0914 treatment were chosen for analysis of immunogenicity. **(C)** Volcano plots depicting change in MHC Class I peptide presentation; select peptides chosen for analysis of immunogenicity are highlighted. **(D)** Charts list average normalized peptide abundance ± SEM for peptides chosen for analysis of immunogenicity (marked with •) as well as for other peptides identified from the same source protein.

## Notes

### Summary of Updates

Manuscript has been extensively revised and new data added

